# Predicting global herbicide resistance hotspots using a 30-year-old database and machine-learning techniques

**DOI:** 10.1101/2023.10.09.561477

**Authors:** Neil Brocklehurst, Chun Liu

## Abstract

The evolution of herbicide resistance in weeds is a problem affecting both food production and ecosystems. Numerous factors affect selection towards herbicide resistance, making it difficult to anticipate where, under what circumstances, and under what timeframe, herbicide resistance is likely to appear. Using the International Herbicide-Resistant Weed Database to provide data on locations and situations where resistance has occurred, we trained models to predict where resistance is most likely in future. Validation of the global models with historical data found a prediction accuracy of up to 78%, while for well-sampled regions, such as Australia, the model correctly predicted more than 95% of instance of resistance and sensitivity. Applying the models to predict instances of resistance over the next decade, future hotspots were detected in North and South America and Australia. Species such as *Conyza canadensis*, *Eleusine indica*, and *Lactuca serriola* are expected to show substantial increases in the number of resistance occurrences. The results highlight the potential of machine-learning approaches in predicting future resistance hotspots and urge more efforts in resistance monitoring and reporting to enable improved predictions. Future work incorporating dimensions such as weed traits, phylogeny, herbicide chemistry, and farming practices could improve the predictive power of the models.

## Introduction

Herbicides provide a simple and usually effective tool for weed management in agriculture, but their use also selects for resistant phenotypes in weed populations. The evolution of herbicide resistance in weeds is a problem affecting both food production and ecosystems [1,2], but it can be delayed, and its spread slowed, with suitable preventative methods based on an understanding of its evolution. Numerous factors affect selection towards herbicide resistance, including weed traits (ecological and biological) [3–6], herbicide mode of action [1,7,8], and agricultural practices [9–11]. This complexity makes it difficult to anticipate where, under what circumstances, and under what timeframe, herbicide resistance is likely to appear. Modelling approaches have provided an avenue for identifying and predicting situations in which resistance may evolve, including mathematical models of the phenomena affecting resistance [12–15], models of population genetics [16–18], and individual-based models addressing the interactions between weed life cycle, spatial distribution, multiple species and agricultural practices [19–21].

The study of herbicide resistance has an advantage over that of fungicide and insecticide resistance, in that there exists a continuously updated global database cases of weed species exhibiting resistance, the herbicides to which they are resistant, and the countries and situations in which resistance is found: the International Herbicide-Resistant Weed Database ([22], http://www.weedscience.org), hereafter the IHRWD. At the time of downloading (20th April 2022), 1639 cases of weed resistance were recorded in the database from between 1957 and 2021 (a case representing resistance in a weed species to a particular mode of action in a particular country/state). This includes 310 weed species in 71 countries, with resistance to 165 active ingredients. The instances of herbicide resistance may be reported by any researcher, and assessed according to a consistent set of criteria to ensure that the data points represent reliable reports. While it is acknowledged that the survey likely under-represents the actual occurrence of herbicide resistance [23], the presence of a unified framework for reporting resistance in a consistent manner provides an opportunity for data-driven approaches for studying and predicting resistance. Similar global databases updated by the research community as a whole (as opposed to a single team or project) exist in other branches of Life Science and are used not only as references and search engines for data points, but also as entire datasets in analyses of broad-scale patterns, acknowledging that all such databases are incomplete and unevenly sampled, but sufficiently broadly sampled to represent global patterns, e.g iNaturalist (conservation and ecology; www.inaturalist.org[e.g. 24-26]), Treebase (phylogenetics; www.treebase.org [e.g. 27,28]), Paleobiology Database (palaeontology; paleobiodb.org [e.g. 29-32]). The IHRWD’s data, while used in historical discussions of resistance [23], has not been leveraged for global scale analysis.

Here we train machine learning models using resistance data from the IHRWD to predict future resistance hotspots globally across countries, weed species, situations (the setting e.g. crop plant, in which the herbicide was applied) and herbicides. Machine learning models have so far been employed at a limited scale in studying herbicide resistance, focussed on individual species within single countries/states [33–36]. The IHRWD provides an opportunity to predict resistance landscapes at a global scale. Our approach validates the concept of using machine learning to predict resistance and identifies areas where additional data and improved screening could improve the reliability of predictions. The models and hotspots presented here should not necessarily be considered as a definitive set of predictions of weed resistance, rather as an exploration of the untapped potential of an existing dataset, and a discussion of potential avenues for future research to produce improved predictions.

## Materials and Methods

### Formation of the Dataset of Herbicide Resistance and Sensitivity

Cases of resistance were downloaded from the International Herbicide-Resistant Weed Database ([22]; www.weedscience.org) on the 20^th^ April 2022. This dataset contains 1639 cases of weed resistance, recorded between 1957 and 2021 (a case representing resistance in a weed species to a particular mode of action in a particular country/state). This includes 310 weed species in 71 countries, with resistance to 165 active ingredients. It should be noted that each case, as recorded in the database, contains multiple herbicides and multiple situations. When these are separated, 8456 unique occurrences of resistance are recorded. Actives from HRAC MOA Group 0 were deleted, as these represent actives of unknown or inconsistent mode of action rather than a distinct group, leaving 8242 occurrences.

Training a model for predicting weed resistance requires not only occurrences of observed weed resistance, but also occurrences of sensitivity i.e., a herbicide is used in a particular situation in a particular country where a particular weed species is present, but no resistant populations have been observed. The sensitive instances were inferred from 1) the geographic ranges of the weeds, drawn from the CABI Invasive species compendium (https://www.cabi.org/isc/), 2) the countries in which crops are grown, identified IndexMundi (https://www.indexmundi.com/), which sources production data from the United States Department of Agriculture (USDA), and World Integrated Trade Solution (https://wits.worldbank.org/), which provides data on crop exports, 3) the situations in which weed species are competitive and 4) the situations and countries in which herbicides have been used, derived from the Pesticide Properties DataBase (PPDB), developed by the Agriculture & Environment Research Unit at the University of Hertfordshire (http://sitem.herts.ac.uk/aeru/ppdb/en/index.htm). If a particular situation is present in a particular country in which a particular herbicide is used, and a competitive weed is present in that country, but no resistant populations have been recorded, it is assumed for the purposes of this study that this represents an instance of sensitivity. Of course, this assumption may not be valid due to the incompleteness of the dataset. Assessment of how this may affect the results is described below.

With data from these sources added, the training dataset produced contained 8242 instances of resistance and 149404 instances of sensitivity. A final issue in dataset formation that needed to be considered before analysis is the fact that the situations are on occasion classified to different levels in the IHRWD. For example, in some cases the situation is defined as “Cereals”, while in others the cereal is specified e.g. “Wheat”. To address this, two different datasets were used in testing. The first used broader situation categories (where necessary, crop types are subsumed into larger categories) or narrow (occurrences with broader categories were deleted).

### Predictor variables

The predictor variables taken from the IHRWD are the weed species, country, situation, herbicide active and MOA. For predicting the future, a predictor variable representing time also needs to be included. Ideally this should be based on how long the active ingredient has been applied in that country but since, as discussed above, such data is not available globally, it is based on length of time since the active ingredient was introduced (treating all actives as if they were released globally simultaneously). This data was obtained from the PPDB. For the training dataset, the time variable for each occurrence is the year the active was introduced subtracted from the current year (2022).

### Testing dataset

The testing dataset includes all occurrences where resistance has not yet been observed (all those counted as sensitive in the training dataset 1957-2021), but with the time variable increased to reflect the fact that 10 years will have passed (2022-2031). Occurrences representing use of active ingredients in countries where they are currently banned, according to the Pan International Consolidated List of Banned Pesticides (https://pan-international.org/pan-international-consolidated-list-of-banned-pesticides/), were deleted from the testing datasets.

### Model Generation and Validation

All machine learning analyses were carried out in the programming language R 4.1.3 [38]. Four different approaches were used to create the models, each from a different class of algorithm: Random Forest (a Decision tree approach) [39,40] implemented in R using functions from the package randomForest [41]; Naïve Bayes (a Bayesian probabilistic approach) implemented using the R package e1071 [42]; Generalised Linear Model (a fexible generalization of ordinary linear regression) [43] implemented in base R; and Support-Vector Machines (classes are separated by a multidimensional hyperplane) [44] implemented using the e1071 package. Stacked models of all four were also tested.

To validate the models and tune the parameters, one would normally randomly subset the training dataset into pseudo-training and pseudo-test datasets and assess how accurately the model predicts the known target variable, and which parameters for the model produced the most accurate predictions e.g. K-fold cross validation. However, since this analysis involves an element of time in the prediction, a slightly different approach was used. The pseudo-training dataset was based on resistance cases before 2017. The models built by the pseudo-training dataset were then used to predict which of the occurrences would show resistance between 2017 and 2021. The performance of the models was assessed using two metrics: Probability of Correct Classification metric (PCC) [45,46], and area under the Receiver Operating Characteristic curve (AUC-ROC) [47,48] (See Supplementary Methods for details of calculation). In the event, both metrics were highly consistent (Table 1), so only the PCC is discussed. In total 2115 models were tested, with five different modelling approaches and various parameters varied and tuned (for full details of the models and parameters see Supplementary Methods).

**Table 1.** Details of the models used in analysis of weed resistance, and statistics regarding their accuracy. AUC-ROC: area under the Receiver Operating Characteristic curve. TPR: True positive rate. TNR: True negative rate. PCC: Probability of correct classification.

|  | Model | Data cutoff | Crop categories | AUC-ROC | TPR | TNR | PCC | TP/TP+FP<br>(Occurrences predicted to evolve resistance that do) | TN/TN+FN<br>(Occurrences predicted to remain sensitive that do) |
| --- | --- | --- | --- | --- | --- | --- | --- | --- | --- |
| Analysis 1 | RF | 75 | Broad | 0.71 | 0.45 | 0.96 | 0.71 | 92% | 64% |
| Analysis 2 | RF | 0 | Narrow | 0.64 | 0.34 | 0.94 | 0.64 | 85% | 59% |
| Analysis 3 | NB | 0 | Broad | 0.53 | 0.74 | 0.32 | 0.53 | 52% | 55% |
| Analysis 4 | NB | 250 | Broad | 0.90 | 0.74 | 0.82 | 0.78 | 80% | 76% |
| Best analysis of Australian dataset | NB | 3 | Broad | 0.95 | 1 | 0.9 | 0.95 | 91% | 100% |
| Best analysis of Australian dataset with abundance as predictor | NB | 3 | Broad | 0.97 | 1 | 0.94 | 0.97 | 93% | 100% |

It should be noted here that an “occurrence”, as used in this paper, represents a unique combination of weed species, country, herbicide active and situation. The possible occurrences are based on the geographic ranges of weeds and the countries in which crops are grown and herbicides are used and assigned as Resistant or Sensitive based on the cases in the IHRWD. It is important to remember when interpreting the results that these are not equivalent to cases of resistance. In a sense, what is being predicted is new database entries. For example, if a particular weed species is not predicted to show numerous novel resistance occurrences over the next 10 years, this does not mean that this weed is not expected to show numerous future cases of resistance, but that cases will not frequently occur in novel situations or countries or to novel herbicides.

### The influence of heterogeneous sampling

The frequency and intensity of resistance screening varies between countries, potentially affecting the frequency of observing cases of resistance [37]. Two proxies were employed to assess how this may have affected the quality of the dataset, and the machine learning predictions. The first assesses the intensity of research in each country, represented by the number of researchers from each country registered with the IHRWD (data taken from the website on 14^th^ September 2022). This may not necessarily be a direct indicator of the frequency of screening: cases reported are not necessarily reported by the teams who found them, or even by researchers in the same country. However, this proxy should provide an insight into the distribution of scientists interested in weed resistance globally, itself indicating to what extent each country prioritises herbicide resistance research. The second is a statistical proxy, assessing the quality of sampling in each country using the metric of coverage, an estimate of the percentage of resistance occurrences in the true population represented by resistant species included in the observed population. This is estimated using Good’s u, a statistic designed in computer science and later adopted in palaeontology and ecology [49–51]. The correlation between the resistance proxies and the absolute number of resistance occurrences was calculated using Pearson’s correlation coefficient, calculated in R.

To show how less heterogenous sampling allows the production of more accurate machine learning models, a more localised analysis was carried out using Australia as a case study. Models were created and validated as described above, replacing Country as a predictor variable with State. The Australian dataset was also used to test an example of how a weedy trait may perform as a predictor: population density of each species within each state. This data was derived from the CABI database as a four-state character: 0: few; 1: localised; 2: present; 3: widespread. This character was included as a predictor variable in creating models from the Australian dataset.

## Results

### Validation of the predictive power of machine learning

The 2115 models predicted resistance occurrences between 2017 and 2021 with accuracies ranging from 0.36 to 0.78, as measured by the PCC. Choosing the best model and set of parameters to use is not simply a matter of choosing the one with the highest overall accuracy; rather it is a compromise between the true positive rate, the true negative rate, and data inclusivity. The models discussed below for predicting resistance hotspots are mostly “cautious” with regards to predicting resistance (Table 1); the occurrences predicted to evolve resistance may be considered reliable predictions, but not a complete list. However, for practical risk assessment for agriculture, worst-case predictions may be more desirable.

Four different models were selected to form the basis of global predictions of resistance up to 2031, based on their performance in the validation (See Table 1 for details of data cutoff, crop treatment and accuracy, see Supplementary Methods and Table S2 for full details of the parameters). Hereafter these are referred to:

– Analysis 1: this model is highly conservative: predictions of resistance may be considered extremely but at the cost of failure to predict many of the resistant occurrences. This analysis also suffers from amalgamating crop categories and not being able to provide predictions for countries and species that did not meet the data cutoff.
– Analysis 2: Similar to analysis 1, this is conservative with regards to predicting resistance. It is less accurate than analysis 1 but includes more data and does not amalgamate crop categories.
– Analysis 3: This was selected as both being inclusive of all data and being less conservative with regards to predictions of resistance: As such predictions for this model may be considered a “worst-case scenario”. This is the least accurate of all four models.
– Analysis 4 is the most accurate model. There is a similar level of accuracy in predictions of resistance and sensitivity. However, like analysis 1 it also suffers from amalgamation of crop categories and has a more severe data cutoff; only seven weed species and five countries meet the cutoff. As such it is not useful for identifying novel future hotspots, since only current hotspots are tested.

When the Australian dataset was subjected to the same validation process, there was again considerable variation in accuracy, as measured by PCC values, but the highest values obtained, 0.95, exceeds any of the models produced from the global dataset. This best-performing model had a true positive rate (correct prediction of resistant occurrences) of 100% and a true negative rate of 90% (Table 1). Such levels of accuracy imply that machine learning has great potential to predict resistance, but that uneven levels of resistance screening between countries limit its accuracy when applied to a global dataset (further analysis of this below).

### Importance of Predictors

Both Random Forest models allow variable importance to be calculated as the mean decrease in GINI index [52] (Table 2). In both analysis 1 and 2, the most important predictor variables are species and country, and the least important is the mode of action (MOA).

**Table 2.** – Importance of predictors (mean decrease in GINI index) calculated for Analyses 1 and 2 (models produced using Random Forest)

| Predictor | Analysis 1 | Analysis 2 |
| --- | --- | --- |
| Weed species | 4125.09 | 11042.28 |
| Country | 4036.44 | 8131.39 |
| Active | 1276.91 | 3716.11 |
| Mode of Action | 892.92 | 2393.66 |
| Situation | 1770.08 | 5596.67 |
| Years since active introduced | 1277.39 | 3553.51 |

### Predictions of Resistance Hotspots

*By Country –* The USA, Canada and Australia are current hotspots of resistance (Fig 1, Fig S4), between them representing more than half of all cases reported in the IHRWD (154 in Australia, 120 in Canada, 584 in the United States), and all analyses predict new occurrences of resistance to continue to appear frequently over the next 10 years. All four analyses suggest the United States will have the highest number of resistance occurrences between 2022 and 2031, and in analyses 1 and 4 (the most accurate), it is the country with the highest proportion of tested occurrences predicted to evolve resistance (11% in Analysis 1, 10% in analysis 4). One should, however, interpret this result with caution; these three countries also account for among the largest numbers of weed scientists active on the IHRWD, and have among the best resistance coverage (Fig 2).

**Figure 1.**
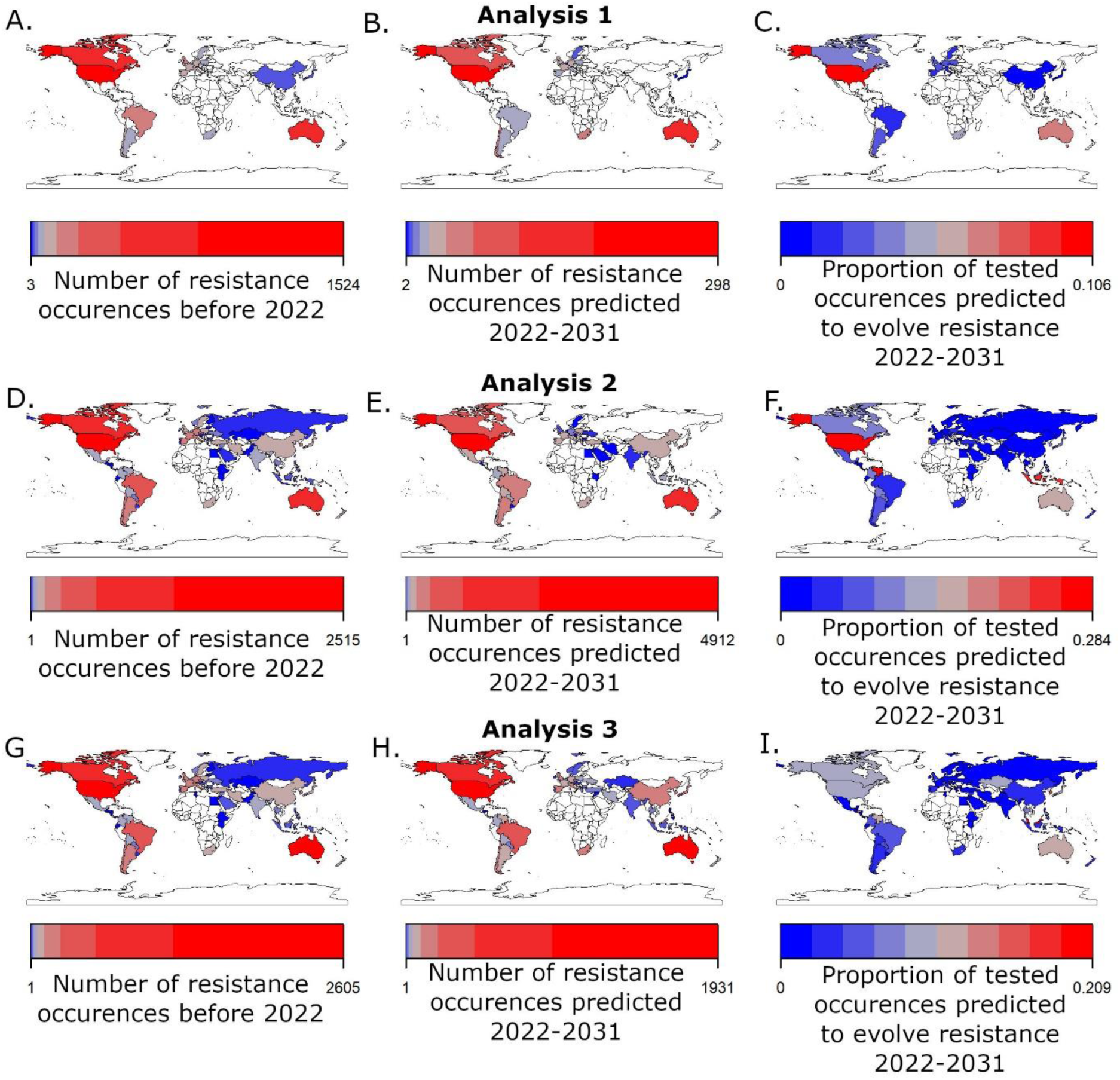
– A-C) Analysis 1; D-E) Analysis 2; G-I) Analysis 3; A, D, G) Number of resistance occurrences observed before 2022, taken from the IHRWD, after data cutoffs applied; B, E, H) Number of resistance occurrences predicted between 2022 and 2031 by the machine learning analyses; C, F, I) Proportion of tested occurrences predicted to evolve resistance between 2022 and 2031 by the machine learning analyses. Results from Analysis 4 presented in Fig S4

**Figure 2.**
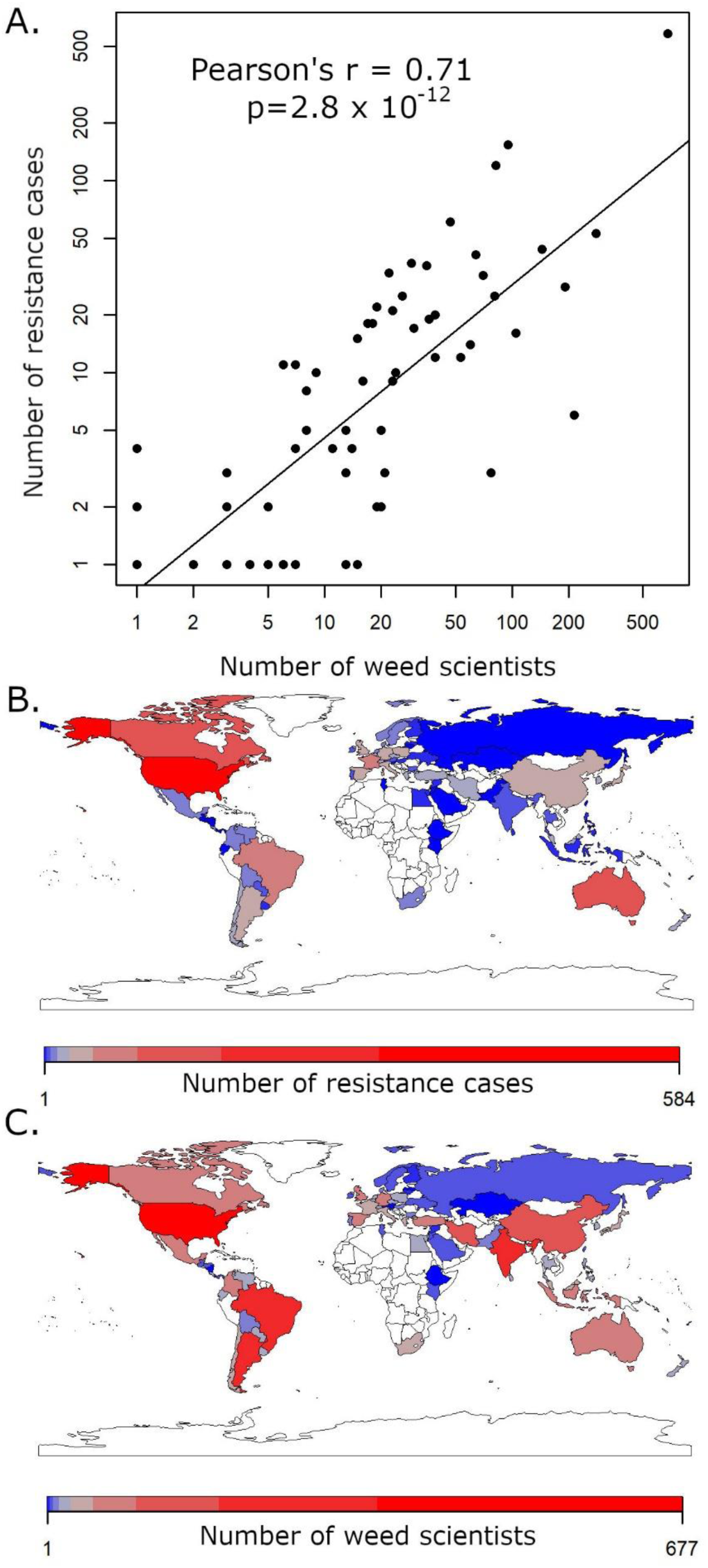
– A) Scattergraph showing the relationship between number of active weed scientists in each country (proxy for sampling intensity) and the number of resistance cases recorded in the IHRWD. C) number of resistance cases recorded in each country; D) number of weed scientists active in each country

South and Central America are also predicted to be foci of future resistance hotspots, although again caution should be taken when interpreting this result, with South American countries accounting for a disproportionate number of active weed scientists (Fig 2D). Analysis 2 and 3 (those with no data cutoff) highlight Venezuela and Costa Rica as areas where a high proportion of the tested occurrences are predicted to show resistance. Analysis 2, the more accurate of the two, predicts in Venezuela 27% of occurrences will evolve resistance, and 23% in Costa Rica (Fig 1d-f). These countries are not tested by analysis 1 due to the data cutoff, but that analysis does highlight Chile as a future resistance hotspot (Fig 1c). Analyses 2 and 3 also indicate Malaysia, South Korea and Japan as areas where a high proportion of the tested occurrences are predicted to show resistance (Analysis 2 predicts in Malaysia 9% of tested occurrences will show resistance, and 8% in Japan and South Korea) (Fig 1f,i).

*By Weed Species –* The predictions for which other weeds will continue to show or develop numerous occurrences of resistance vary from analysis to analysis. Analyses 1, 2 and 4 (the most accurate analyses) predict *Lolium rigidum* (annual ryegrass) and *Lolim perenne multiflorum* (Italian ryegrass), two weed species with the large numbers of resistance cases recorded in the present (*L. perenne multiflorum* has 71 cases recorded in the IHRWD, more than any other weed, with resistance to eight MOAs recorded in 14 countries) will continue to evolve numerous novel cases of resistance in future. Thirty seven percent of tested occurrences of *L. rigidum* in analysis 1, 19% in analysis 2 and 16% in Analysis 4, were predicted to show resistance; the proportions in *L. perenne multiflorum* are 21, 12 and 9% respectively (Fig 3, Fig S5).

**Figure 3.**
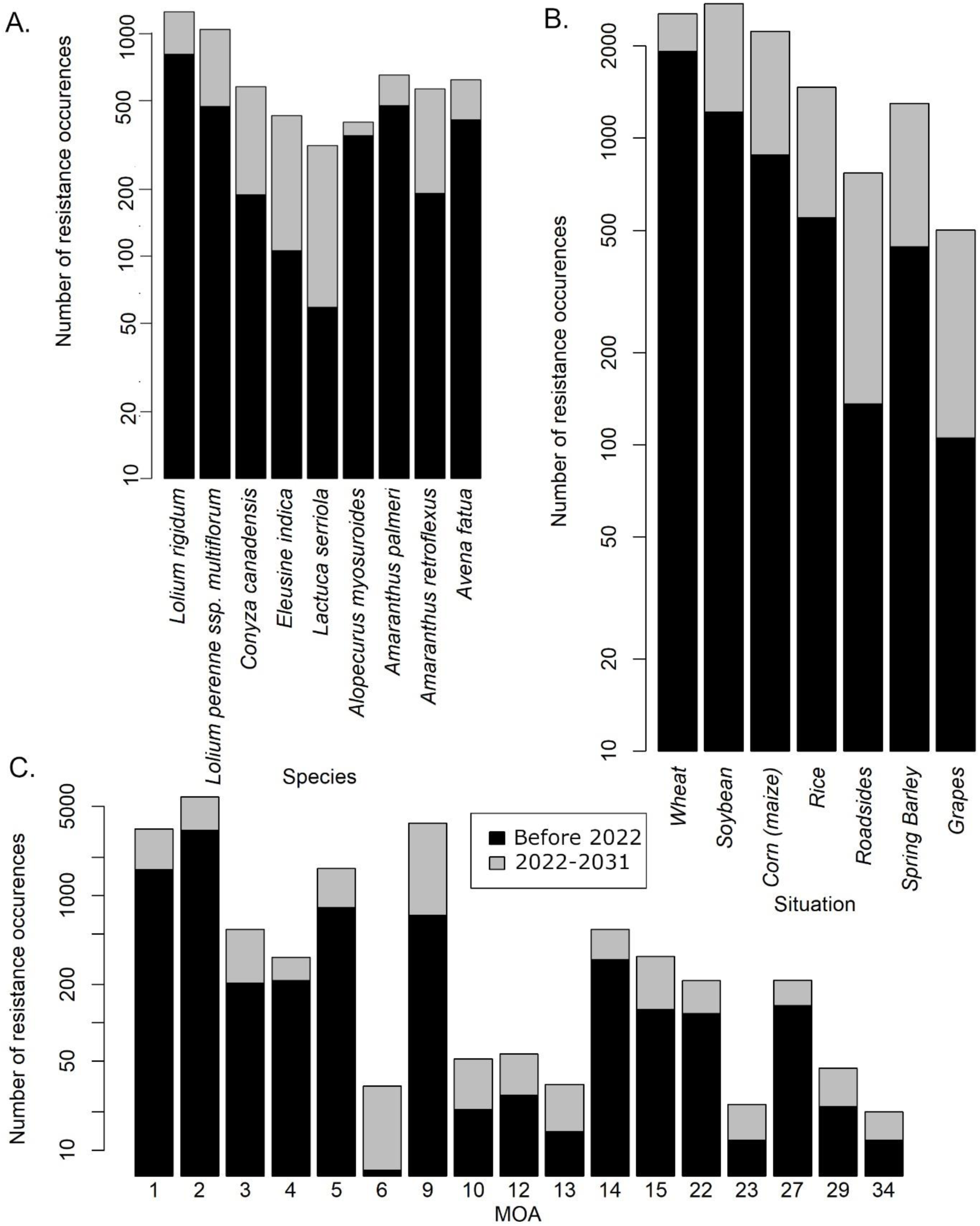
– The number resistance occurrences from 1957 until 2021, derived from the IHRWD (black) and from 2022-2031 in A) selected weed species; B) Selected situations; C) HRAC mode of action classifications.

*Conyza canadensis* (Canadian horseweed) is predicted by all three analyses in which it is included to show a substantial increase in the number of resistance occurrences (it not included in Analysis 4 due to not meeting the data cutoff) (Fig 3, Fig S5, S6). Analysis 1 and 2 both found that of tested occurrences involving *C. canadensis*, 12% were predicted to evolve resistance, while in analysis 3 the proportion was 10%, similar to or substantially more than other weeds with more frequent cases of resistance recorded in the present e.g., in Analysis 2, *Amaranthus palmeri* is predicted to evolve resistance in 11% of tested occurrences; *Amaranthus retroflexus* in 10%; *Avena fatua* in 9%; *Alopecurus myosuroides* in 2% (Fig 3, Fig S5). The majority of the predicted resistance occurrences in *C. canadensis* are expected to occur in the United States, but Indonesia, Hungary, Portugal and Japan are predicted to show a higher frequency than in the present (Fig S7).

Analyses 2 and 3, where no data cut-offs were applied, both predict a high proportion of the tested occurrences of *Eleusine indica* and *Lactuca serriola* to show resistance. *E. indica* (Indian goosegrass) does not currently show particularly frequent evolution of resistance (37 cases in the IHRWD) but resistance to eight different MOAs has been found, potentially indicating a high intrinsic ability to evolve resistance. In analysis 2 (the more accurate of the two), 11% of tested occurrences of *E. indica* are expected to show resistance (Fig 3, Fig S5), and in analysis 3 (the worst-case scenario) it is 12%.

*By Situation –* Analyses 1 and 3 (which use broad crop categories) predict numerous occurrences of resistance in beans, with analysis 2 (narrow crop categories) singling out soybeans (Fig 3, Fig S5, S8). Analyses 1,2 and 4 (those with the greatest accuracy) predict numerous occurrences of resistance in corn in future. Resistance in soybean and corn situations is most frequently found in the USA and Canada in the present, and these countries are predicted to remain hotspots (Fig S9).

There are some situations that are less frequently sites of resistance in the present day, yet predicted to become hotspots of resistance. Analyses 1 and 3 (which use broad crop categories) predict numerous occurrences of resistance in Cereal settings, but analysis 2 (narrow crop categories) singles out spring barley rather than wheat as the future hotspot. Of the tested occurrences with spring barley as the situation, 855 (18%) were predicted to evolve resistance over the next 10 years (with the United States in particular showing substantial increases in the number of occurrences), compared to 623 (2%) in wheat (Fig 3, Fig S5).

*By herbicide* – The largest numbers of resistance cases in the IHWRD are found in herbicides of the modes of action of HRAC groups 1 (Inhibition of acetyl CoA carboxylase, ACCase), 2 (Inhibition of ALS), 5 (Inhibition of Photosystem II) and 9 (Inhibition of Enolpyruvyl Shikimate Phosphate Synthase [glyphosate only]). Analyses 1, 2 and 3 predict that over the next 10 years, groups 1, 2 and 9 will continue to produce large numbers of resistance occurrences (Fig 5, Fig S10. Analyses 2, and 3 (no data cutoff) also highlight those of HRAC groups 3 (Inhibition of Microtubule Assembly) and 5 (Inhibition of Photosystem II) (Fig 3, Fig S5). Analysis 4, with its high data cutoff, only tests herbicides from four modes of action (1, 2, 5 and 9), with 5 and 9 predicted to have the largest number and proportion of tested occurrences showing resistance. Results regarding other modes of action are highly inconsistent between the four sets of analyses, but two consistent results that the herbicide actives glyphosate and fenoxaprop-p-ethyl (a group 1 herbicide) show large numbers of resistance cases now and are predicted under all four analyses to show large numbers of resistance occurrences in future.

### Impact of variation in screening frequency

The variation in screening frequency and intensity by country, and how it may affect frequency of identifying cases of resistance, may be demonstrated by the correlation between proxies for sampling effort and the number of resistance cases reported in each country: The number of weed science researchers in each country and the coverage of resistant species in that country (Good’s u). Both proxies show a strong and significant correlation with the number of resistance cases in each country (Fig 2), implying that the amount of resistance observed in a country is heavily linked to the quality of sampling and the intensity of weed-resistance research from each country. The Pearson’s R value for the correlation between number of weed scientists from each country and number of resistanc cases is 0.71 (p= 2.8 x 10^-12^), and between the coverage of reistant weed species (Good’s U) and number of resistance cases it is 0.77 (p = 7.1 x 10^-15^)

## Discussion

### The database’s predictive potential

The IHRWD, although incomplete, contains a broad sampling of resistance cases from across the world, and so provides a promising source of data for global analyses of resistance patterns. Nevertheless, there are issues with using the data uncritically to predict resistance occurrences, and many such issues are highlighted by the analyses herein.

As described above, the quality of sampling (the frequency and intensity of screening for resistance) potentially impacts the relative frequencies of resistance observations [37].

The direct impact of sampling quality on the frequency of resistance observations is demonstrated by the strong and highly significant correlations between the number of resistance cases in each country and the two sampling proxies, in particular the number of weed scientists from each country. This not only affects our impressions of the current status of resistance but also the accuracy of predictive models; larger numbers of resistance cases in a particular country in the training dataset will bias towards predicting resistance in that country, and it is difficult to separate instances where such large numbers are genuine or due to sampling inconsistencies.

The fact that the subset of the IHRWD representing resistance cases within a single well-screened country (Australia) produced a more accurate predictive models than the global dataset further demonstrates the impact of sampling bias on the global models. An analysis within a single country should reduce the impact of such sampling heterogeneity, as it is expected that the frequency of resistance screening will be more consistent between states than between countries. The Australian models are not only more accurate than the global models, but have a high absolute accuracy of predictions, with a probability of correct classification of 95%. This demonstrates that the data within the IHRWD does have predictive power, although at present the global dataset are not so accurate as those using a local subset. Incorporation of sampling correction and extrapolation methods commonly applied to ecological data [50–53] could improve accuracy, but building individual models for countries with frequent resistance screening may be a more promising avenue.

### The Importance of Predictors

Variation in the importance of predictor variables in a model is not only useful for understanding and evaluating the predictions made, but, being derived from the training dataset, can provide insights into how important the various factors are in driving resistance.

That country is an important variable is not surprising. As discussed above, a large element of the frequency of cases of resistance may be driven by resistance screening practices, which vary by country. The country also combines the signals of the geographic range of weed species, the range in which crops are grown, and agronomic practices. The high importance of weed species perhaps indicates a great influence of the intrinsic ability of certain weeds to evolve resistance. Traits such as seed production and dispersal mode determine how rapidly resistant populations grow and spread [3,5]. Pollination mode and dioecy of plants influence the genetic diversity within populations [3,6]. Such traits vary by plant and will influence the frequency with which resistant populations appear. Future models could incorporate the traits directly as predictors, not only potentially improving the accuracy of the models but also informing on the relative importance of traits as drivers of increased resistance risk. As an example, a predictor representing population density within states as a five-state discrete character was added to the Australian dataset. The validation produced a small increase in the accuracy of predictions (Table 1).

The low variable importance of the mode of action, in particular relative to that of the specific active ingredient, might be related to the fact that these represent nested variables; each active belongs to a broader MOA category, and so the Active variable represents a more specific indicator of herbicide chemistry. However, the reduced importance of mode of action appears to be a more recent phenomenon; training the models on data from earlier years shows Active and MOA have similar importance, with the importance of Active increasing during the 1990s (Fig S1,2). The timing of this follows shortly on an increase in the frequency of introduction of herbicides (Fig S3), as well as an increase in the use of modes of action developed post-1970 [54], potentially driving increased competition between actives for baseline effectiveness and resistance-breaking.

### Predictions of resistance hotspots

*Prediction of resistance hotspots by weed species* – Two results that are consistent across analyses are the continued high frequency of resistance in multiple species of *Lolium*, and the increased number of resistance occurrences in *Conyza canadensis*. *C. canadensis* has been the subject of many past studies of weed resistance, notably as being the first weed in which glyphosate resistance was reported [55]. Although it has a relatively large number of resistance cases reported in the IHRWD (66), resistance to only five MOAs is reported. More than half of cases (37) are to glyphosate alone, and the majority are from the United States. In a recent assessment of resistance risk in the European and Mediterranean Plant Protection Organization area, Moss et al. [56] classified *Conyza* as “Medium risk”. Nevertheless, *C. canadensis* has also been highlighted as being intrinsically prone to evolving resistance to photosystem I inhibitors (HRAC group 22) [23,57], as well as having a particularly strong tendency to evolve resistance to multiple modes of action.

On the other hand, the prediction of continued frequent resistance in *Lolium* is not surprising. This weed species has been discussed as being particularly prone to resistance. *L. rigidum* in particular, the species predicted to show the most frequent occurrences of resistance, is noted as representing the first reported case of resistance to acetolactate synthase-inhibiting herbicides [58]. It has a high degree of genetic variability and appears to be particularly prone to stacking of resistance [23].

*Amaranthus* is another highly problematic weed genus with numerous cases of resistance recorded the IHRWD: to 18 modes of action in 11 species across 21 countries. Herbicide resistance in *Amaranthus* is considered a serious threat to cropping systems, and the genus show an exceptional tendency to evolve resistance, with a rapid growth rate, exceptional seed production, and dioecy forcing outcrossing [6]. *A. tuberculatus*, the common waterhemp, was found by the most recent WSSA surveys to be the most common and most troublesome weed species in soy, wheat and corn fields [59,60]. Unfortunately, despite the clear necessity for accurate predictions of the resistance future in *Amaranthus*, the models are unable to provide consistent results. Analysis 1 does not predict frequent new occurrences of resistance evolving in any *Amaranthus* species; the largest number is in *A. retroflexus* (common amaranth), with 32 occurrences (fewer than weeds such as *Stellaria media*, *Avena fatua* and *Avena sterilis*, all of which have fewer cases of resistance recorded in the IHRWD), representing fewer than 2% of tested occurrences. Analysis 2 predicts a large number of new occurrences in *Amaranthus retroflexus*: 368 (more comparable with such species as *Conyza canadensis* and *Kochia scoparia*), representing 10% of tested occurrences. Analysis 3, however, predicts numerous occurrences of resistance in *A. tuberculatus* (23% of tested occurrences) and *A. palmeri* (palmer amaranth; 14%), but less frequent occurrences in *A. retroflexus* (6%). Analysis 4 (the most accurate analysis, but only testing six weed species) predicted *A. palmeri* would evolve more occurrences of resistance than any other tested weed, both in terms of absolute numbers (31) and proportion of tested occurrences (56%). *Amaranthus* may be a case where a more detailed look at more local results, particularly in the USA where it is especially problematic, could provide a more reliable assessment than the global analysis.

The large number of resistance occurrences predicted in *L. serriola* (prickly lettuce) is surprising, as recorded instances of resistance are much less frequent (only seven cases in the IHRWD; the species did not make the data cutoff to be included in analyses 1 and 4). This species is less frequently a target for herbicides as its early germination reduces competition with some crops [61]. Nevertheless, despite this low number of cases and being a relatively infrequent target of herbicides, resistance to three different MOAs has evolved, and it represented of the first documented case confirmed to be target-site resistance to ALS-inhibiting herbicides [62], potentially indicating a high intrinsic tendency to rapidly evolve resistance when it is exposed to herbicides. When one examines the herbicides to which *L. serriola* has evolved resistance in the IHRWD, the length of time between the introduction of herbicide and earliest case of resistance to that herbicide has a median of 2.5 years; only *Alopecurus myosuroides* has lower (Fig S12). The rapidity with which it evolves resistance may explain why it is predicted to evolve resistance in 20% of tested occurrences by 2031 in analysis 2.

*Prediction of resistance hotspots by situation* – The situations in which herbicide resistance are most frequently found in the present are wheat, corn, soybeans and rice, which have accounted for the majority of herbicide usage over the longest time [23]. Roadsides and orchards are also sites where resistance is commonly found, being sites of frequent application to produce bare ground [23]. However, the large increases in frequency of resistance predicted by models in spring barley and grapes are less consistent with past reviews of the database.

Barley has in recent years been highlighted as a more competitive crop than wheat [63,64] particularly against *Lolium* and *Alopecurus* whose autumn emergence also renders spring cropping an effective grass weed control strategy [65,66]. While its competitiveness with weeds makes it a favoured choice for use in crop rotations and intercropping, its increased use in this context does make it a more frequent setting for herbicide use and therefore resistance evolution.

Analyses 1 and 3 also predict frequent occurrences in fruit, but Analysis 2 singles out grapes rather than any fruit grown in orchards (Fig 3, Fig S5). This, however, may be an artefact of the way situations are grouped. Under the narrow crop categories orchards (which may represent multiple different fruits) are deleted. This does highlight the categorisation of crops within the IHRWD as an issue that can affect the results, due to the inconsistency in the level of category that the situation is defined at. When this is considered, the fact that grapes are predicted to be a potential resistance hotspot is less surprising. After removing the categories of “Orchards” and “Fruit”, grapes are the most frequent situation for weed resistance in the fruit category. The IHRWD records 39 cases of weed resistance with grapes as the situation in 13 different countries, most frequently in the United States, South Africa, Australia, New Zealand, and France (Fig S9). 17% of tested occurrences with grapes as the situation are predicted to show resistance by 2031 (Fig 3, Fig S5). Like orchards, vineyards employ extensive and repeated use of herbicides throughout the year to keep the under-vine ground bare [67,68]. The long cropping cycle prevents the use of crop rotation to manage weeds [69], and “hilling” (building up of soil around the base of the vines for winter protection) has been shown to increase weed density, potentially bringing seeds to the surface and promoting germination [70].

*Prediction of resistance hotspots by herbicide* – The different models produce inconsistent predictions regarding which herbicides and herbicide MOA groups are likely to become problematic. However, three MOA groups were highlighted by all four analyses: ACCase inhibiting herbicides (Group 1), ALS inhibiting herbicides (Group 2) and ESPS inhibiting herbicides (Group 9; glyphosate only).

Glyphosate has been a commercially important herbicide, effective on almost all weeds and able to treat massive areas due to the development of herbicide-tolerant crops. It was also thought to be highly unlikely to select for resistance, due to its low soil activity and the limited number of target-site mutations conferring resistance [5,23]. Resistant weeds were not discovered until 22 years after its introduction. However, it is worth noting that glyphosate use was not so prevalent until the introduction of tolerant crops in 1996, prior to which glyphosate was used in a similar manner and frequency to non-selective herbicides like diquat and paraquat [71]. Resistance to these herbicides appeared 25 and 18 years after their respective introductions, so the 22-year interval observed in glyphosate is not so unusual. The perception of limited number of target site mutations and fitness cost in wild populations has also been re-evaluated [72,73]. Due to its now-widespread use on many weed species, resistance has become extremely common: 342 cases are reported in the IHRWD in 30 countries and 55 species. That it is predicted to show increases in resistance occurrences in future is to be expected, and of the tested occurrences involving glyphosate, 55% were predicted to show resistance according to Analysis 2 (Fig 3, Fig S5).

Group 2 herbicides represent the most frequent cases of resistance in the present day (676 cases in the IHRWD), although this is potentially due in part to the large number of actives within the group (at the time of writing there are 57, more than any other group). They are the most widely and frequently used herbicides [23]. However, there are factors beyond the numerical aspects that make these herbicides intrinsically more prone to selecting for resistance. Resistance to ALS-inhibiting herbicides was found less than five years after their introduction [62], and the frequency of cases has increased more rapidly than other widely-used herbicide groups [7]. Suggestions for factors that may drive this increased tendency to resistance include target-site resistance being controlled by single dominant nuclear encoded genes [74,75], flexibility of the target site allowing multiple different amino acid substitutions to confer resistance [7], and a lack of significant fitness cost to resistance [76,77]. However, it should be noted that the large absolute number of resistance cases predicted over the next 10 years does not represent a large proportion of tested occurrences in any analyses (Fig 3, Fig S10), implying that use over a wide area and variety of situations is still a large factor in the frequency of resistance.

Group 1 herbicides are applied to grass weeds (family Poaceae) and have been widely employed in cereals-growing situations, leading to numerous infestations of resistant grass weeds [23]. The fact that *Lolium* is predicted to show further increases in the frequency of resistant occurrences, and that cereals are predicted to be a frequent situation for resistant occurrences in future, may well be driving the substantial increases in the frequency of resistance predicted for this group. Fenoxaprop-p-ethyl has 121 recorded cases of resistance in 33 countries reported in the IHRWD, making resistance to this active more widespread than any other. However, the large number of resistance occurrences predicted in the future appear to be more due to the large number of situations and countries in which it is used and the large number of target species providing more opportunities for resistance; although the absolute number is large, the proportion of occurrences with fenoxaprop-p-ethyl as active predicted by analysis 2 to show resistance is only 5%, and for group 1 as a whole it is 7% (Fig 3, Fig S5).

## Conclusions

The International Herbicide-Resistant Weed Database represents a valuable resource allowing a variety of data-driven approaches for analysis at a global scale. From a modelling perspective, perfect dataset rarely exists, and it is not necessarily best practice to defer modelling efforts until all desired data are available. Rather, model development can specify data gaps and guide further data collection activities, triggering new modelling cycle for improved prediction [78]. The work presented here highlights the value of machine learning models in the prediction of weed resistance and meanwhile advocates more efforts in resistance monitoring and reporting, at both local and global scales, and preferably also recording sensitive cases. More detailed and finely tuned models, could represent useful tools for identifying priorities for increased resistance screening, monitoring and proactive actions and targets for selective herbicide research and development. Broad-scale analyses of weed resistance is an area of considerable potential.

## Acknowledgements

We would like to thank V. Grimm and D. Kaundun for helpful comments that greatly improved the manuscript. J. Downes, I. Heap, S.-J. Hutchings, F. Kon, G. Le Goupil, G. Namou Jablak, W. Plumb, R Shukla also provided useful discussion. *Thanks to reviewers to be added*

## References

1. Powles, S. B., & Yu, Q. (2010). Evolution in action: plants resistant to herbicides. Annual review of plant biology, 61, 317–347.

2. Mortensen, D. A., Egan, J. F., Maxwell, B. D., Ryan, M. R., & Smith, R. G. (2012). Navigating a critical juncture for sustainable weed management. BioScience, 62, 75–84.

3. Jasieniuk, M., Brûlé-Babel, A. L., & Morrison, I. N. (1996). The evolution and genetics of herbicide resistance in weeds. Weed science, 44, 176–193.

4. Christoffers, M. J. (1999). Genetic aspects of herbicide-resistant weed management. Weed Technology, 13, 647–652.

5. Nandula, V. K. (2010). Herbicide resistance: Definitions and concepts. Glyphosate resistance in crops and weeds: History, development, and management, 35–43.

6. Tranel, P. J. (2021). Herbicide resistance in Amaranthus tuberculatus. Pest Management Science, 77, 43–54.

7. Tranel, P. J., & Wright, T. R. (2002). Resistance of weeds to ALS-inhibiting herbicides: what have we learned? Weed Science, 50, 700–712.

8. Moss, S. (2017). Herbicide resistance in weeds. Weed research: Expanding horizons, 181–214.

9. Diggle, A. J., Neve, P. B., & Smith, F. P. (2003). Herbicides used in combination can reduce the probability of herbicide resistance in finite weed populations. Weed Research, 43, 371–382.

10. Neve, P., & Powles, S. (2005). High survival frequencies at low herbicide use rates in populations of Lolium rigidum result in rapid evolution of herbicide resistance. Heredity, 95, 485–492.

11. Beckie, H. J., & Reboud, X. (2009). Selecting for weed resistance: herbicide rotation and mixture. Weed Technology, 23, 363–370.

12. Gressel, J., & Segel, L. A. (1978). The paucity of plants evolving genetic resistance to herbicides: possible reasons and implications. Journal of Theoretical Biology, 75, 349–371.

13. Gressel, J., & Segel, L. A. (1990). Modelling the effectiveness of herbicide rotations and mixtures as strategies to delay or preclude resistance. Weed Technology, 4, 186–198.

14. Birch, C. P. D., & Shaw, M. W. (1997). When can reduced doses and pesticide mixtures delay the build-up of pesticide resistance? A mathematical model. Journal of applied ecology, 1032–1042.

15. Gustafson DI (2008) Sustainable use of glyphosate in North American cropping systems. Pest Manag Sci 64:409–416

16. Maxwell, B. D., Roush, M. L., & Radosevich, S. R. (1990). Predicting the evolution and dynamics of herbicide resistance in weed populations. Weed technology, 4(1), 2–13.

17. Jasieniuk, M., & Maxwell, B. D. (1994). Populations genetics and the evolution of herbicide resistance in weeds. Phytoprotection, 75, 25–35.

18. Diggle, A. J., & Neve, P. (2001). The population dynamics and genetics of herbicide resistance—a modeling approach. In Herbicide resistance and world grains (pp. 61–99). CRC Press.

19. Liu, C., Bridges, M. E., Kaundun, S. S., Glasgow, L., Owen, M. D., & Neve, P. (2017). A generalised individual-based algorithm for modelling the evolution of quantitative herbicide resistance in arable weed populations. Pest management science, 73, 462–474.

20. Somerville GJ, Powles SB, Walsh MJ, Renton M (2017). How do spatial heterogeneity and dispersal in weed population models affect predictions of herbicide resistance evolution? Ecological Modelling, 362, 37–53.

21. Thornby D, Werth J, Hereward J, Keenan M, Chauhan BS (2018). Herbicide resistance evolution can be tamed by diversity in irrigated Australian cotton: a multi-species, multi-herbicide modelling approach. Pest Management Science, 74, 2363–75.

22. Heap, I. The International Herbicide-Resistant Weed Database. Online. April 20, 2022

23. Heap, I. (2014). Global perspective of herbicide-resistant weeds. Pest management science, 70(9), 1306–1315.

24. Calzada Preston, C. E., & Pruett-Jones, S. (2021). The number and distribution of introduced and naturalized parrots. Diversity, 13, 412.

25. Rosa, R. M., Cavallari, D. C., & Salvador, R. B. (2022). iNaturalist as a tool in the study of tropical molluscs. Plos one, 17, e0268048.

26. Cull, B. (2022). Monitoring trends in distribution and seasonality of medically important ticks in North America using online crowdsourced records from iNaturalist. Insects, 13, 404.

27. Blum, M. G., & François, O. (2006). Which random processes describe the tree of life? A large-scale study of phylogenetic tree imbalance. Systematic Biology, 55, 685–691.

28. Le, S. Q., Lartillot, N., & Gascuel, O. (2008). Phylogenetic mixture models for proteins. Philosophical Transactions of the Royal Society B: Biological Sciences, 363, 3965–3976.

29. Alroy, J. (2008). Dynamics of origination and extinction in the marine fossil record. Proceedings of the National Academy of Sciences, 105, 11536–11542.

30. Foote, M. (2007). Extinction and quiescence in marine animal genera. Paleobiology, 33, 261–272.

31. Dunhill, A. M., Hannisdal, B., Brocklehurst, N., & Benton, M. J. (2018). On formation–based sampling proxies and why they should not be used to correct the fossil record. Palaeontology, 61, 119–132.

32. Brocklehurst, N., Kammerer, C. F., & Benson, R. J. (2020). The origin of tetrapod herbivory: effects on local plant diversity. Proceedings of the Royal Society B, 287, 20200124.

33. Stratonovitch, P., Storkey, J., & Semenov, M. A. (2012). A process-based approach to modelling impacts of climate change on the damage niche of an agricultural weed. Global Change Biology, 18, 2071–2080.

34. Storkey, J., Stratonovitch, P., Chapman, D. S., Vidotto, F., & Semenov, M. A. (2014). A process-based approach to predicting the effect of climate change on the distribution of an invasive allergenic plant in Europe. PloS one, 9, e88156.

35. Hicks, H., Lambert, J., Pywell, R., Hulmes, L., Hulmes, S., Walker, K., … & Freckleton, R. P. (2021). Characterizing the environmental drivers of the abundance and distribution of Alopecurus myosuroides on a national scale. Pest Management Science, 77, 2726–2736.

36. Evans, J. A., Tranel, P. J., Hager, A. G., Schutte, B., Wu, C., Chatham, L. A., & Davis, A. S. (2016). Managing the evolution of herbicide resistance. Pest management science, 72, 74–80.

37. Brodersen, K. H., Ong, C. S., Stephan, K. E., & Buhmann, J. M. (2010, August). The balanced accuracy and its posterior distribution. In 2010 20th international conference on pattern recognition (pp. 3121–3124). IEEE.

38. Suzen, M. (2020). A simple and interpretable performance measure for a binary classifier. Memo’s Island

39. Fawcett, T. (2006). An introduction to ROC analysis. Pattern Recognition Letters, 27, 861–874.

40. Han, J., Zhu, L., Kulldorff, M., Hostovich, S., Stinchcomb, D.G., Tatalovich, Z., Lewis, D.R. and Feuer, E.J., 2016. Using Gini coefficient to determining optimal cluster reporting sizes for spatial scan statistics. International journal of health geographics, 15(1), pp.1–11.

41. Good, I. J. (1953). The population frequencies of species and the estimation of population parameters. Biometrika, 40, 237–264.

42. Peterson, M. A., Collavo, A., Ovejero, R., Shivrain, V., & Walsh, M. J. (2018). The challenge of herbicide resistance around the world: a current summary. Pest management science, 74, 2246–2259.

43. Alroy, J. (2010). Geographical, environmental and intrinsic biotic controls on Phanerozoic marine diversification. Palaeontology, 53(6), 1211–1235.

44. Chao, A., & Jost, L. (2012). Coverage-based rarefaction and extrapolation: standardizing samples by completeness rather than size. Ecology, 93, 2533–2547.

45. Hsieh, T. C., Ma, K. H., & Chao, A. (2016). iNEXT: an R package for rarefaction and extrapolation of species diversity (Hill numbers). Methods in Ecology and Evolution, 7(12), 1451–1456.

46. Osteen, C.D. and Fernandez-Cornejo, J., 2016. Herbicide use trends: a backgrounder. Choices, 31(4), pp.1–7.

47. VanGessel, M. J. (2001). Glyphosate-resistant horseweed from Delaware. Weed Science, 49, 703–705.

48. Moss, S., Ulber, L., & den Hoed, I. (2019). A herbicide resistance risk matrix. Crop protection, 115, 13–19.

49. Weaver, S., Downs, M., & Neufeld, B. (2004). Response of paraquat-resistant and-susceptible horseweed (Conyza canadensis) to diquat, linuron, and oxyfluorfen. Weed science, 52, 549–553.

50. Heap, I., & Knight, R. (1986). The occurrence of herbicide cross-resistance in a population of annual ryegrass, Lolium rigidum, resistant to diclofop-methyl. Australian Journal of Agricultural Research, 37, 149–156.

51. Van Wychen L (2020) 2020 Survey of the most common and troublesome weeds in grass crops, pasture, and turf in the United States and Canada. Weed Science Society of America National Weed Survey Dataset.

52. Van Wychen L (2019) 2019 Survey of the most common and troublesome weeds in broadleaf crops, fruits & vegetables in the United States and Canada. Weed Science Society of America National Weed Survey Dataset

53. Preston, C., Stone, L. M., Rieger, M. A., & Baker, J. (2006). Multiple effects of a naturally occurring proline to threonine substitution within acetolactate synthase in two herbicide-resistant populations of Lactuca serriola. Pesticide Biochemistry and Physiology, 84, 227–235.

54. Mallory-Smith, C. A., Thill, D. C., & Dial, M. J. (1990). Identification of sulfonylurea herbicide-resistant prickly lettuce (Lactuca serriola). Weed technology, 4, 163–168.

55. Doll, H., Holm, U., and Søgaard, B. (1995). Effect of crip density on competition by wheat and barley with Agrostemma githago and other weeds. Weed Research 35, 391–396

56. Molla, A., & Sharaiha, R. K. Competition and resource unitization in mixed cropping of barley and durum wheat under different moisture stress levels. World Journal of Agricultural Sciences, 6, 713–719

57. Bertholdsson, N. O. (2005). Early vigour and allelopathy–two useful traits for enhanced barley and wheat competitiveness against weeds. Weed Research, 45, 94–102.

58. Moss, S., & Lutman, P. (2013). Black-grass: the potential of non-chemical control. Harpenden: Rothamsted Research.

59. Dastgheib, F., & Frampton, C. (2000). Weed management practices in apple orchards and vineyards in the South Island of New Zealand. New Zealand Journal of Crop and Horticultural Science, 28, 53–58.

60. Nordblom, T., Penfold, C., Whitelaw-Weckert, M., Norton, M., Howie, J., & Hutchings, T. (2021). Financial comparisons of under-vine management systems in four South Australian vineyard districts. Australian Journal of Agricultural and Resource Economics, 65, 246–263.

61. Hanson, B., Wright, S., Sosnoskie, L., Fischer, A., Jasieniuk, M., Roncoroni, J., … & Al-Khatib, K. (2014). Maintaining long-term management: herbicide-resistant weeds challenge some signature cropping systems. California Agriculture, 68, 142–152.

62. Jiang, L., Koch, T., Dami, I., & Doohan, D. (2008). The effect of herbicides and cultural practices on weed communities in vineyards: an Ohio survey. Weed Technology, 22, 91–96.

63. Duke, S. O. (2018). The history and current status of glyphosate. Pest Management Science, 74, 1027–1034.

64. Shaner, D. L., Lindenmeyer, R. B., & Ostlie, M. H. (2012). What have the mechanisms of resistance to glyphosate taught us? Pest Management Science, 68, 3–9.

65. Vila-Aiub, M.M., Yu, Q. and Powles, S.B., 2019. Do plants pay a fitness cost to be resistant to glyphosate?. New Phytologist, 223(2), pp.532–547.

66. Sebastian, S. A., Fader, G. M., Ulrich, J. F., Forney, D. R., & Chaleff, R. S. (1989). Semidominant soybean mutation for resistance to sulfonylurea herbicides. Crop Science, 29, 1403–1408.

67. Hart, S. E., Saunders, J. W., & Penner, D. (1993). Semidominant nature of monogenic sulfonylurea herbicide resistance in sugarbeet (*Beta vulgaris*). Weed Science, 41, 317–324.

68. Holt, J. S., & Thill, D. C. (1994). Growth and productivity of resistant plants. 299–316. SB Powles and JAM Holtum. Herbicide Resistance in Plants: Biology and Biochemistry. Boca Raton, FL Lewis.

69. Blackshaw, R. E., Kanashiro, D., Moloney, M. M., & Crosby, W. L. (1994). Growth, yield and quality of canola expressing resistance to acetolactate synthase inhibiting herbicides. Canadian journal of plant science, 74(4), 745–751.

70. Schmolke A, Thorbek P, DeAngelis DL, Grimm V. Ecological models supporting environmental decision making: a strategy for the future. Trends in Ecology & Evolution. 2010; 25:479–86.

71. R Core Team (2022). R: A language and environment for statistical computing. R Foundation for Statistical Computing, Vienna, Austria.

72. Ho, T. K. (1998). The random subspace method for constructing decision forests. IEEE transactions on pattern analysis and machine intelligence, 20(8), 832–844.

73. Breiman, L. (2001). Random forests. Machine learning, 45(1), 5–32.

74. Liaw, A., & Wiener, M. (2002). Classification and regression by randomForest. R news, 2(3), 18–22.

75. Meyer, D., Dimitriadou, E., Hornik, K., Weingessel, A., Leisch, F., Chang, C. C., … & Meyer, M. D. (2019). Package ‘e1071’. The R Journal.

76. Nelder, J. A. Wedderburn, (1972).: Generalized Linear Model”. Journal of the Royal Statistical Society. Series A (General), 135(3).

77. Cortes, C., & Vapnik, V. (1995). Support-vector networks. Machine learning, 20, 273–297.

78. Anthony, M., & Holden, S. B. (1998). Cross-validation for binary classification by real-valued functions: theoretical analysis. In Proceedings of the eleventh annual conference on Computational learning theory (pp. 218–229).

