## Supplementary Methods for "Predicting global herbicide resistance hotspots using a 30-year-old database and machine-learning techniques": Supplementary Data 26.R

```
######
###
### Two functions (The first for the datasets used for
### Random Forest,SVM and Naive Bayes, the second for the datasets used in 
### glm only) that calculate the proportion of resistant and sensitive
### occurences each year from 1957 (the first resistance occurence in the 
### database) until the selected year, fit a second order polynomial
### curve to this data, then use it to predict the expected proportion in a
### later year. 
### Both functions work in the same way (the 
### only difference is the fact that the first expects resistance to be 
### scored as "A" for resistant and "B" for sensitive, the latter as "1" and 
### "0" respectively. The arguments in both are:
###
### ML.data - the training dataset, as created in supplementary data 25,26 and
###           27
###
### data.year - the year at which your data stops (2016 for tuning, 2022 for 
###             analysis)
###
### test.year - the year you wish to predict to (2022 for tuning, 2031 for 
###             analysis
###
######

find.prior<-function(ML.data,data.year,test.year)
{


	years<-1957:data.year#####vector of years

	prop.res<-vector(length=length(years))#########empty vector to store proportion of resistant occurences

	for(i in 1:length(years))#####each year one by one
	{
		test.year<-years[i]####year to test

		year.data<-ML.data[which(ML.data$Active.year<=test.year),] ####delete actives not present uin that year
	
		resist.rows<-which(year.data$Resistance=="A") #####resistant occurences
		too.young.rows<-intersect(resist.rows,which(year.data$Year>test.year))#######find those where resistance was not observed until after 
	
		year.data[too.young.rows,"Resistance"]<-"B"#####Set them as sensitive

		prop.res[i]<-length(which(year.data$Resistance=="A"))/nrow(year.data)####find proportion

	}

	plot(years,prop.res)######plot proportion

	year.data<-data.frame(years,prop.res)   ####fit polynomial model and add line
	line<-lm(prop.res~poly(years,2),data=year.data)
	lines(years,predict(line,year.data))


	year.data.2<-rbind(year.data,c(test.year,NA))######extrapolate line to test year
	predictions<-predict(line,year.data.2)
	return(predictions[length(predictions)])
}


find.prior.glm<-function(ML.data,data.year,test.year)
{


	years<-1957:data.year

	prop.res<-vector(length=length(years))

	for(i in 1:length(years))
	{
		test.year<-years[i]

		year.data<-ML.data[which(ML.data$Active.year<=test.year),]
	
		resist.rows<-which(year.data$Resistance==1)
		too.young.rows<-intersect(resist.rows,which(year.data$Year>test.year))
	
		year.data[too.young.rows,"Resistance"]<-0

		prop.res[i]<-length(which(year.data$Resistance==1))/nrow(year.data)

	}

	plot(years,prop.res)

	year.data<-data.frame(years,prop.res)

	line<-lm(prop.res~poly(years,2),data=year.data)


	lines(years,predict(line,year.data))


	year.data.2<-rbind(year.data,c(test.year,NA))
	predictions<-predict(line,year.data.2)
	return(predictions[length(predictions)])
}
```
