## Supplementary Methods for "Predicting global herbicide resistance hotspots using a 30-year-old database and machine-learning techniques": Supplementary Data 27.R

```
######################
#####
##### Extracting occurences (unique combinations of Species, Country,	
##### Active and Situation) from the cases in the database
#####
######################

weed.data<-read.csv(file.choose())######Read in Supplementary Data 1 (Data downloaded from the resisatcne database)


clean.weed<-matrix(nrow=0,ncol=7)#####empty matrix

for(i in 1:nrow(weed.data))#######Each row one by one
{
	row<-weed.data[i,]######extract row


	######
	###
	###Commas in cells indicate multiple entries. find commas and 
	###create separate entries for each
	###
	######
	
	####MOA
	if(grepl(", ",row[1,5]))####If comma present...
	{
		MOA.entries<-strsplit(row[1,5],split=", ")[[1]] ###Separate into multiple entries...
	}
	else(MOA.entries<-row[1,5])####otherwise keep as one entry
	
	####Same for Actives
	if(grepl(", ",row[1,6]))
	{
		active.entries<-strsplit(row[1,6],split=", ")[[1]]
	}
	else(active.entries<-row[1,6])
	
	####Same for Situations
	if(grepl(", ",row[1,7]))
	{
		crop.entries<-strsplit(row[1,7],split=", ")[[1]]
	}
	else(crop.entries<-row[1,7])

	#########
	####
	#### Create a new entry for each mode of action for each active, 
	#### for each crop
	####
	#########
		
	for(x in 1:length(MOA.entries)) #####each mode of action entry
	{
		for(y in 1:length(active.entries))#### each active entry
		{
			for(z in 1:length(crop.entries))#### each situation entry
			{

				######Create new row
				new.entry<-row
				new.entry[1,5]<-MOA.entries[x]
				new.entry[1,6]<-active.entries[y]
				new.entry[1,7]<-crop.entries[z]
	
				clean.weed<-rbind(clean.weed,new.entry)###attach new row to empty matrix
			}
		}
	}		

}


##################
##################
##################
#
# Weed distribuitions
#
##################
##################
##################


CABI.weed<-read.csv(file.choose(),header=F) #####Read in Supplementary Data 2 (weed distributions from CABI)


weed.countries<-matrix(nrow=length(unique(clean.weed$Species)),######matrix to store distributions
	ncol=length(unique(c(CABI.weed[,1],clean.weed$Country))),data=0)


rownames(weed.countries)<-unique(clean.weed$Species) #####rows are weed species
colnames(weed.countries)<-unique(c(CABI.weed[,1],clean.weed$Country))#####columns are countries

for(i in unique(clean.weed$Species))######each species 1 by 1
{
	weed.dist<-unique(c(clean.weed$Country[which(clean.weed$Species==i)],#####find what countries each weed is recorded in in the database and CABI
		CABI.weed[,1][which(CABI.weed[,2]==i)]))


	weed.countries[i,weed.dist]<-1######set those countries as present (1) in the matrix
}


##################
##################
##################
#
# Crop distribuitions
#
##################
##################
##################

crop.trade<-read.csv(file.choose(),na.strings="",row.name=1)######Read in supplementray data 3 (crop distributions)

situation.countries<-matrix(nrow=length(unique(clean.weed$Situations)), #####Matrix to record distributions
	ncol=length(unique(c(clean.weed$Country,unlist(strsplit(
	crop.trade[which(is.na(crop.trade[,1])==F),1],split=", "))))),data=0)


rownames(situation.countries)<-unique(clean.weed$Situations)#####rows are situations 
colnames(situation.countries)<-sort(unique(c(clean.weed$Country,unlist( #####columns are countries
	strsplit(crop.trade[which(is.na(crop.trade[,1])==F),1],split=", ")))))


for(i in unique(clean.weed$Situations))######Each situation 1 by 1
{
	sit.dist<-unique(c(clean.weed$Country[which(clean.weed$Situations==i)],######What countries each situation is recorded in in the database and the file
		strsplit(crop.trade[i,1],split=", ")[[1]]))
	sit.dist<-sit.dist[is.na(sit.dist)==F]
	
	situation.countries[i,sit.dist]<-1######set those countries as present (1) in the matrix

}


situation.countries[c("Railways","Roadsides",#####set these four as present in all countries
	"Fencelines","Around Buildings"),]<-1


##################
##################
##################
#
# Herbicides situations of use
#
##################
##################
##################


herbicides<-read.csv(file.choose(),row.names=1)#######read in supplementary data 4 (situations in which herbicides are used

herbicide.situations<-matrix(nrow=length(unique(clean.weed$Actives)),#####Matrix to record situations of use
	ncol=length(unique(clean.weed$Situations)),data=0)

rownames(herbicide.situations)<-unique(clean.weed$Actives)#####rows are actives
colnames(herbicide.situations)<-unique(clean.weed$Situations)####columns are situations


for(i in unique(clean.weed$Actives))######each active one by 1
{
	herb.sit<-unique(c(clean.weed$Situations[which(clean.weed$Actives==i)],####What situations each active is recorded in in the database and the file
		strsplit(herbicides[i,2],split=", ")[[1]]))
	herb.sit<-herb.sit[is.na(herb.sit)==F]
	
	herbicide.situations[i,herb.sit]<-1######set those situations as used (1) in the matrix


}


##################
##################
##################
#
# Herbicides countries of use
#
##################
##################
##################


herbicide.countries<-matrix(nrow=length(unique(clean.weed$Actives)),######Matrix to store distributions
	ncol=length(unique(clean.weed$Country)),data=0)

rownames(herbicide.countries)<-unique(clean.weed$Actives)#####rows are actives
colnames(herbicide.countries)<-unique(clean.weed$Country)#####columns are countries


for(i in unique(clean.weed$Actives))#####each active one by one
{
	herb.dist<-unique(clean.weed$Country[which(clean.weed$Actives==i)])######find countries where active is recorded in the database

	
	herbicide.countries[i,herb.dist]<-1######set those countries as present (1) in the matrix


}


##################
##################
##################
#
# Situations where weeds are competitive
#
##################
##################
##################


weed.situations<-matrix(nrow=length(unique(clean.weed$Species)),######Matrix to store weed/situation combos
	ncol=length(unique(clean.weed$Situations)),data=0)

rownames(weed.situations)<-unique(clean.weed$Species)#####rows are weed species
colnames(weed.situations)<-unique(clean.weed$Situations)####colums are situations


for(i in unique(clean.weed$Species))######each species one by one
{
	herb.sit<-unique(clean.weed$Situations[which(clean.weed$Species==i)])######find situations where weed is recorded in the database
	weed.situations[i,herb.sit]<-1

}


##################
##################
##################
#
# Create full dataset with sensistives
#
##################
##################
##################

Resistance<-rep("A",nrow(clean.weed))####Addd new column to databse showiung resisatant/sensitive (at the momment, everything A [Resistant]
test.weed<-cbind(clean.weed,Resistance)


############################
#### Convert MOA to HRAC number (quicker to work with shorter strings)
############################

herbicides<-unique(test.weed$Actives)#### Unique actives

MOA<-read.csv(file.choose(),head=F,row.name=1)######Read in Supplementary Data 5 (MOAs of Actives)
MOA<-as.vector(MOA[,1])#######Convert number to character
MOA<-as.character(MOA)
names(MOA)<-herbicides#####assigne herbicides as names

for(i in test.weed$Actives)
{
	test.weed[which(test.weed$Actives==i),"MOAs"]<-MOA[i]####go through database and change MOA ciolumn to single character
}


############################
#### Add sensitives
############################

for(i in unique(test.weed$Species))#######each species one by one
{
	weed.range<-colnames(weed.countries)[which(weed.countries[i,]==1)]#####which countries is weed found in

	weed.crop<-colnames(weed.situations)[which(weed.situations[i,]==1)]#####which crops is weed competitive in

	for(j in weed.crop)######each competitive situation one by one
	{
		crop.range<-colnames(situation.countries)[ #######which countries is that situation present in 
			which(situation.countries[j,]==1)]
	
		crop.actives<-rownames(herbicide.situations)[ #####which herbicides are used in that situation 
			which(herbicide.situations[,j]==1)]

		for(k in crop.actives)###### each of the used herbicides one by one
		{
			herbicide.range<-colnames(herbicide.countries)[######what countries is that herbicide used in
				which(herbicide.countries[k,]==1)]

			test.countries<-intersect(crop.range,intersect(weed.range,########intersection between the weed,crop and herbicide ranges
				herbicide.range))

			resist<-intersect(which(test.weed$Country%in%test.countries),#######which of the intersections is found in the resistance database
				intersect(which(test.weed$Species==i),
				intersect(which(test.weed$Situations==j),
				which(test.weed$Actives==k))))


			sensitive.countries<-test.countries[#####those that arn't are sensitive countries
				which(test.countries%in%
				test.weed[resist,"Country"]==F)]

			if(length(sensitive.countries)!=0)#####if there are any sensitive counrties.....
			{
				#####Create new rows for sensitive combinations
				new.rows<-matrix(nrow=length(sensitive.countries),#####Empty matrix withone row for each sensitive country
					ncol=ncol(test.weed))
				colnames(new.rows)<-colnames(test.weed)
				new.rows<-as.data.frame(new.rows)####convert to data frame
	
				#####Fill columns
	
				new.rows$Year<-1957#####Current year (just to find something to fill it with; this column is not relevant for senistives except it needs to be older than 2016)
				new.rows$Species<-i
				new.rows$Country<-sensitive.countries
				new.rows$Actives<-k
				new.rows$Situations<-j
				new.rows$Resistance<-"B"######Resistance column set as "B" (sensitive)
				new.rows$MOAs<-MOA[k]


				test.weed<-rbind(test.weed,new.rows)#####add new rows


			}
		}
	}
}

test.weed<-test.weed[which(test.weed$MOAs!="0"),]#######delete rows with group 0 herbicides	
	
```
