## Supplementary Methods for "Predicting global herbicide resistance hotspots using a 30-year-old database and machine-learning techniques": Supplementary Data 28.R

```
library(e1071)
library(randomForest)
library(glmnet)
library(klaR)
library(vioplot)
library(rworldmap)


#########
###
### prepping data
###
#########

load(file.choose()) #####Read in supplementary data 6

years<-read.csv(file.choose(),header=F,row.name=1)#####Read in supplementary data 7
crop.categories<-read.csv(file.choose(),header=F)#####Read in supplementary data 8 for analyses 1 and 4, supplementary data 9 for analyses 2 and 3

Active.year<-years[test.weed$Active,]#### addd the year of active to the dataset
test.weed<-cbind(test.weed,Active.year)


#######convert the situations based on broad or narrow crop categories

for(i in 1:nrow(crop.categories))
{
	if(crop.categories[i,2]!="")
	{
		if(crop.categories[i,2]=="Delete")
		{
			rows.to.delete<-which(test.weed$Situations==
				crop.categories[i,1])
			test.weed<-test.weed[-rows.to.delete,]
		}
		else
		{
			test.weed$Situations[which(test.weed$Situations==
				crop.categories[i,1])]<-crop.categories[i,2]
		}
	}
}


data.cutoff<-0######set data cutoff (0 for analysis 2, 75 for analysis 1)


weed.species<-sort(unique(test.weed$Species))######take weed species

weeds.to.check<-names(which( #######find those which meet data cutoff
	table(test.weed$Species[which(test.weed$Resistance=="A")])>=data.cutoff))


countries<-sort(unique(test.weed$Country))######take countries

countries.to.check<-names(which( #######find those which meet data cutoff 
	table(test.weed$Country[which(test.weed$Resistance=="A")])>=data.cutoff))


herbicides<-sort(unique(test.weed$Actives))######take actives

herbicides.to.check<-names(which( #######find those which meet data cutoff 
	table(test.weed$Actives[which(test.weed$Resistance=="A")])>=data.cutoff))


test.year<-2021
predict.year<-2031


ML.data<-test.weed[,c("Year","Species","Country","MOAs","Actives","Situations","Resistance","Active.year")]#####Creeate full dataset

ML.data<-ML.data[which(ML.data$Species%in%weeds.to.check),]
ML.data<-ML.data[which(ML.data$Country%in%countries.to.check),]
ML.data<-ML.data[which(ML.data$Actives%in%herbicides.to.check),]


year.data<-ML.data[which(ML.data$Active.year<=test.year),]

resist.rows<-which(year.data$Resistance=="A")
too.young.rows<-intersect(resist.rows,which(year.data$Year>test.year))

year.data[too.young.rows,"Resistance"]<-"B"

time<-test.year-year.data$Active.year
train<-cbind(year.data,time)


###############Analysis 1 and 2 (random forest)

######For analysis 2 only: current priors
weights.0<-1/length(which(year.data$Resistance=="A"))
weights.1<-1/length(which(year.data$Resistance=="B"))

######For analysis 1 only (predictive priors
prior<-find.prior(ML.data,test.year,predict.year)
weights.0<-(1-prior)
weights.1<-prior


######For both
prior.weights<-vector(length=nrow(year.data))######set priors vector
prior.weights[which(year.data$Resistance=="A")]<-weights.0
prior.weights[which(year.data$Resistance=="B")]<-weights.1


#######Create model
###Analysis 1
model.rf <- randomForest(y=as.factor(train$Resistance),######create model
	x=train[,c("Situations","Species","Country","MOAs","Actives","time")],
	mtry=3,ntree=1500,weights=prior.weights)

###Analysis 2
model.rf <- randomForest(y=as.factor(train$Resistance),######create model
	x=train[,c("Situations","Species","Country","MOAs","Actives","time")],
	mtry=1,ntree=1500,weights=prior.weights)


######create test dataset (any occurneces where ressiatnce has not yet evolved, with time advanced to meet the test year
test.rows<-which(ML.data$Resistance=="B")
time<-predict.year-ML.data$Active.year[test.rows]
test<-cbind(ML.data[test.rows,],time)
test<-test[is.na(time)==F,]


######Remove banned herbicides

bans<-read.csv(file.choose(), header=T,row.name=1,check.names=F)#####Read in Supplementary Data 10 (herbicide ban data)

rows.removed<-vector(length=0)####empty vector to store rows removed
for(i in 1:nrow(test))######each occurence one by one
{
	herbicide<-train[i,"Actives"]####The active
	country<-train[i,"Country"]#####The country

	if(herbicide %in% rownames(bans) && country %in%colnames(bans))#####if they are in the ban matrix
	{		
		if(is.na(bans[herbicide,country])==F)#####If banned
		{
			rows.removed<-c(rows.removed,i)####addd occurence to list to be deleted
		}
	}
}


test<-test[-rows.removed,]#####remove banned rows


######Make prediction
prediction<-predict(model.rf,test)
future.resist<-test[which(prediction=="A"),]#####which are predicted to show resistance?


###############Analysis 3 and 4 (Naive Bayes)

#####Analysis 3
model.nb<-NaiveBayes(x=train[,c("Situations","Country","Actives","MOAs",
	"Species","time")],grouping=as.factor(train$Resistance),
	 usekernel=T,fL=2,prior=c(0.5,0.5))
#####Analysis 4
model.nb<-NaiveBayes(x=train[,c("Situations","Country","Actives","MOAs",
	"Species","time")],grouping=as.factor(train$Resistance),
	 usekernel=T,fL=0,prior=c(0.5,0.5))


######create test dataset (any occurneces where ressiatnce has not yet evolved, with time advanced to meet the test year
test.rows<-which(ML.data$Resistance=="B")
time<-predict.year-ML.data$Active.year[test.rows]
test<-cbind(ML.data[test.rows,],time)
test<-test[is.na(time)==F,]


######Remove banned herbicides (only needed for analysis 3)
bans<-read.csv(file.choose(), header=T,row.name=1,check.names=F)#####Read in Supplementary Data 10 (herbicide ban data)

rows.removed<-vector(length=0)
for(i in 1:nrow(test))
{
	herbicide<-train[i,"Actives"]
	country<-train[i,"Country"]

	if(herbicide %in% rownames(bans) && country %in%colnames(bans))
	{		
		if(is.na(bans[herbicide,country])==F)
		{
			rows.removed<-c(rows.removed,i)
		}
	}
}

test<-test[-rows.removed,]


####Make predictiion
prediction<-predict(model.nb,test)$class
future.resist<-test[which(prediction=="A"),]


#########
###
### Resistance by country
###
#########


future.country<-table(future.resist$Country)#########count of predicted occurences by country
potential.country<-table(test$Country)#######count of tested occurences
past.country<-table(ML.data$Country[########Count of database occurences
	which(ML.data$Resistance=="A")])

prop.country<-future.country/potential.country#####proportioon of tested occurences predicted to show resistance


########
###
###Future resistance in species
###
########


future.spec<-table(future.resist$Species)#####count of predicted occurences by species
past.spec<-table(ML.data$Species[######count of database occurences
	which(ML.data$Resistance=="A")])
potential.spec<-table(test$Species)#######count of tested occurences

prop.spec<-future.spec/potential.spec#####proportion of tested occurences predicted to show resistance


#####
##
## resistance change in situation
##
#####

future.sit<-table(future.resist$Situations)#####count of predicted occurences by situation
past.sit<-table(ML.data$Situations[which(ML.data$Resistance=="A")])######count of database occurences
potential.sit<-table(test$Situations)#######count of tested occurences


prop.sit<-future.sit/potential.sit#####proportion of tested occurences predicted to show resistance


#####
##
## resistance change in Actives
##
#####

future.act<-table(future.resist$Actives)#####count of predicted occurences by active
past.act<-table(ML.data$Actives[which(ML.data$Resistance=="A")])######count of database occurences
potential.act<-table(test$Actives)#######count of tested occurences

prop.sit<-future.sit/potential.sit#####proportion of tested occurences predicted to show resistance


#####
##
## resistance change by MOA
##
#####
MOAs<-read.csv(file.choose(),header=F,row.name=1)#######Read in supplementary data 5 (MOAs of actives)

future.MOA<-cbind(future.act,MOAs[names(future.act),])#####count of predicted occurences by MOA
future.MOA<-aggregate(future.MOA[,1], by=list(future.MOA[,2]), FUN=sum)

past.MOA<-cbind(past.act,MOAs[names(past.act),])######count of database occurences
past.MOA<-aggregate(past.MOA[,1], by=list(past.MOA[,2]), FUN=sum)

potential.MOA<-cbind(potential.act,MOAs[names(potential.act),])#######count of tested occurences
potential.MOA<-aggregate(potential.MOA[,1], by=list(potential.MOA[,2]), FUN=sum)


prop.MOA<-future.MOA[,2]/potential.MOA[,2]#####proportion of tested occurences predicted to show resistance
```
