## Supplementary Methods for "Predicting global herbicide resistance hotspots using a 30-year-old database and machine-learning techniques": Supplementary Data 29.R

```
library(e1071)
library(randomForest)
library(glmnet)
library(klaR)
library(vioplot)
library(rworldmap)


find.prior<-function(ML.data,data.year,test.year)
{


	years<-1957:data.year

	prop.res<-vector(length=length(years))

	for(i in 1:length(years))
	{
		test.year<-years[i]

		year.data<-ML.data[which(ML.data$Active.year<=test.year),]
	
		resist.rows<-which(year.data$Resistance=="A")
		too.young.rows<-intersect(resist.rows,which(year.data$Year>test.year))
	
		year.data[too.young.rows,"Resistance"]<-"B"

		prop.res[i]<-length(which(year.data$Resistance=="A"))/nrow(year.data)

	}

	plot(years,prop.res,pch=16,xlab="Year",ylab="Cumulative proportion of resistant occurences")

	year.data<-data.frame(years,prop.res)

	line<-lm(prop.res~poly(years,2),data=year.data)


	lines(years,predict(line,year.data),lwd=3)


	year.data.2<-rbind(year.data,c(test.year,NA))
	predictions<-predict(line,year.data.2)
	return(predictions[length(predictions)])
}


find.prior.glm<-function(ML.data,data.year,test.year)
{


	years<-1957:data.year

	prop.res<-vector(length=length(years))

	for(i in 1:length(years))
	{
		test.year<-years[i]

		year.data<-ML.data[which(ML.data$Active.year<=test.year),]
	
		resist.rows<-which(year.data$Resistance==1)
		too.young.rows<-intersect(resist.rows,which(year.data$Year>test.year))
	
		year.data[too.young.rows,"Resistance"]<-0

		prop.res[i]<-length(which(year.data$Resistance==1))/nrow(year.data)

	}

	plot(years,prop.res)

	year.data<-data.frame(years,prop.res)

	line<-lm(prop.res~poly(years,2),data=year.data)


	lines(years,predict(line,year.data))


	year.data.2<-rbind(year.data,c(test.year,NA))
	predictions<-predict(line,year.data.2)
	return(predictions[length(predictions)])
}


load(file.choose())

years<-read.csv(file.choose(),header=F,row.name=1)
crop.categories<-read.csv(file.choose(),header=F)


Active.year<-years[test.weed$Active,]
test.weed<-cbind(test.weed,Active.year)


for(i in 1:nrow(crop.categories))
{
	if(crop.categories[i,2]!="")
	{
		if(crop.categories[i,2]=="Delete")
		{
			rows.to.delete<-which(test.weed$Situations==
				crop.categories[i,1])
			test.weed<-test.weed[-rows.to.delete,]
		}
		else
		{
			test.weed$Situations[which(test.weed$Situations==
				crop.categories[i,1])]<-crop.categories[i,2]
		}
	}
}


###############Predicting the futures with Random forest
data.cutoff<-0


weed.species<-sort(unique(test.weed$Species))######Extract weed species

weeds.to.check<-names(which( #######put limit on number of occurences of resistance to 
	table(test.weed$Species[which(test.weed$Resistance=="A")])>=data.cutoff))


countries<-sort(unique(test.weed$Country))######Extract weed species

countries.to.check<-names(which( #######put limit on number of occurences of resistance to 
	table(test.weed$Country[which(test.weed$Resistance=="A")])>=data.cutoff))


herbicides<-sort(unique(test.weed$Actives))######Extract weed species

herbicides.to.check<-names(which( #######put limit on number of occurences of resistance to 
	table(test.weed$Actives[which(test.weed$Resistance=="A")])>=data.cutoff))


test.year<-2021
predict.year<-2031


ML.data<-test.weed[,c("Year","Species","Country","MOAs","Actives","Situations","Resistance","Active.year")]#####Creeate full dataset

ML.data<-ML.data[which(ML.data$Species%in%weeds.to.check),]
ML.data<-ML.data[which(ML.data$Country%in%countries.to.check),]
ML.data<-ML.data[which(ML.data$Actives%in%herbicides.to.check),]


year.data<-ML.data[which(ML.data$Active.year<=test.year),]

resist.rows<-which(year.data$Resistance=="A")
too.young.rows<-intersect(resist.rows,which(year.data$Year>test.year))

year.data[too.young.rows,"Resistance"]<-"B"

time<-test.year-year.data$Active.year
train<-cbind(year.data,time)


weights.0<-1/length(which(year.data$Resistance=="A"))
weights.1<-1/length(which(year.data$Resistance=="B"))


prior<-find.prior(ML.data,test.year,predict.year)
weights.0<-(1-prior)
weights.1<-prior


prior.weights<-vector(length=nrow(year.data))
prior.weights[which(year.data$Resistance=="A")]<-weights.0
prior.weights[which(year.data$Resistance=="B")]<-weights.1


model.rf <- randomForest(y=as.factor(train$Resistance),
	x=train[,c("Situations","Species","Country","MOAs","Actives","time")],
	mtry=1,ntree=1500,weights=prior.weights)


test.rows<-which(ML.data$Resistance=="B")
time<-predict.year-ML.data$Active.year[test.rows]
test<-cbind(ML.data[test.rows,],time)
test<-test[is.na(time)==F,]


######herbicide bans

bans<-read.csv(file.choose(), header=T,row.name=1,check.names=F)

rows.removed<-vector(length=0)
for(i in 1:nrow(test))
{
	herbicide<-train[i,"Actives"]
	country<-train[i,"Country"]

	if(herbicide %in% rownames(bans) && country %in%colnames(bans))
	{		
		if(is.na(bans[herbicide,country])==F)
		{
			rows.removed<-c(rows.removed,i)
		}
	}
}


test<-test[-rows.removed,]

prediction<-predict(model.rf,test)


future.resist<-test[which(prediction=="A"),]


###############Predicting the futures with Naive Bayes
data.cutoff<-0


weed.species<-sort(unique(test.weed$Species))######Extract weed species

weeds.to.check<-names(which( #######put limit on number of occurences of resistance to 
	table(test.weed$Species[which(test.weed$Resistance=="A")])>=data.cutoff))


countries<-sort(unique(test.weed$Country))######Extract weed species

countries.to.check<-names(which( #######put limit on number of occurences of resistance to 
	table(test.weed$Country[which(test.weed$Resistance=="A")])>=data.cutoff))


herbicides<-sort(unique(test.weed$Actives))######Extract weed species

herbicides.to.check<-names(which( #######put limit on number of occurences of resistance to 
	table(test.weed$Actives[which(test.weed$Resistance=="A")])>=data.cutoff))


test.year<-2021
predict.year<-2031


ML.data<-test.weed[,c("Year","Species","Country","MOAs","Actives","Situations","Resistance","Active.year")]#####Creeate full dataset

ML.data<-ML.data[which(ML.data$Species%in%weeds.to.check),]
ML.data<-ML.data[which(ML.data$Country%in%countries.to.check),]
ML.data<-ML.data[which(ML.data$Actives%in%herbicides.to.check),]


year.data<-ML.data[which(ML.data$Active.year<=test.year),]

resist.rows<-which(year.data$Resistance==1)
too.young.rows<-intersect(resist.rows,which(year.data$Year>test.year))

year.data[too.young.rows,"Resistance"]<-0

time<-test.year-year.data$Active.year
train<-cbind(year.data,time)


model.nb<-NaiveBayes(x=train[,c("Situations","Country","Actives","MOAs",
	"Species","time")],grouping=as.factor(train$Resistance),
	 usekernel=T,fL=2,prior=c(0.5,0.5))


test.rows<-which(ML.data$Resistance=="B")
time<-predict.year-ML.data$Active.year[test.rows]
test<-cbind(ML.data[test.rows,],time)
test<-test[is.na(time)==F,]


######herbicide bans

bans<-read.csv(file.choose(), header=T,row.name=1,check.names=F)

rows.removed<-vector(length=0)
for(i in 1:nrow(test))
{
	herbicide<-train[i,"Actives"]
	country<-train[i,"Country"]

	if(herbicide %in% rownames(bans) && country %in%colnames(bans))
	{		
		if(is.na(bans[herbicide,country])==F)
		{
			rows.removed<-c(rows.removed,i)
		}
	}
}


test<-test[-rows.removed,]


prediction<-predict(model.nb,test)$class


future.resist<-test[which(prediction=="A"),]


#########
###
### All resistance
###
#########


future.country<-table(future.resist$Country)
potential.country<-table(test$Country)
past.country<-table(ML.data$Country[
	which(ML.data$Resistance=="A")])


future.country[names(past.country)[which(names(past.country)%in%
	names(future.country)==F)]]<-0
future.country<-future.country[names(past.country)]

potential.country[names(past.country)[which(names(past.country)%in%
	names(potential.country)==F)]]<-0
potential.country<-potential.country[names(past.country)]
prop.country<-future.country/potential.country
prop.country<-as.vector(prop.country)
names(prop.country)<-names(future.country)


names(past.country)[names(past.country)%in%countryExData$Country==F]

data(countryExData)
countryExData$Country[15]<-"Bosnia and Herzegovina"
countryExData$Country[21]<-"Central African Republic"
countryExData$Country[26]<-"C�te D'ivoire"
countryExData$Country[28]<-"Democratic Republic of Congo"
countryExData$Country[34]<-"Czech Republic"
countryExData$Country[38]<-"Dominican Republic"
countryExData$Country[145]<-"Vietnam"
countryExData$Country[134]<-"Trinidad and Tobago"


country.codes<-countryExData[which(countryExData$Country%in%names(past.country)),
	c("Country","ISO3V10")]


plot.past.country<-data.frame(past.country[country.codes[,1]],
	country.codes[,2])
names(plot.past.country)<-c("Name","Freq","Code")


resist.map <- joinCountryData2Map( plot.past.country
	,joinCode = "ISO3"
	,nameJoinColumn = "Code")


plot.count.country<-data.frame(as.array(future.country[country.codes[,1]]),
	country.codes[,2])
names(plot.count.country)<-c("Name","Freq","Code")


count.map <- joinCountryData2Map( plot.count.country
	,joinCode = "ISO3"
	,nameJoinColumn = "Code")


plot.future.country<-data.frame(as.array(future.country[country.codes[,1]]/
	potential.country[country.codes[,1]]),country.codes[,2])
names(plot.future.country)<-c("Name","Freq","Code")


future.map <- joinCountryData2Map( plot.future.country
	,joinCode = "ISO3"
	,nameJoinColumn = "Code")


tiff(filename = "Analysis_3_country.tif",width = 20, height = 10, 
	units = "cm",res=300, compression = "lzw")

par(mfrow=c(1,3))
par(mar=c(0,1,0,0.5))

mapCountryData(resist.map,nameColumnToPlot="Freq",
	catMethod="logFixedWidth",colourPalette=c("blue","grey","red"),
	borderCol="black",numCats = 10,mapTitle = "")
mapCountryData(count.map,nameColumnToPlot="Freq",
	catMethod="logFixedWidth",colourPalette=c("blue","grey","red"),
	borderCol="black",numCats = 10,mapTitle = "")
mapCountryData(future.map,nameColumnToPlot="Freq",
	catMethod="fixedWidth",colourPalette=c("blue","grey","red"),
	borderCol="black",numCats = 10,mapTitle = "")
dev.off()


combined<-rbind(past.country,future.country)
colnames(combined)<-names(future.country)
rownames(combined)<-c("Past","Predicted")

tiff(filename = "FigureS3.tif",width = 20, height = 20, 
	units = "cm",res=300, compression = "lzw")

par(mfrow=c(1,2))
par(mar=c(8,5,1,1))
barplot(combined, xlab = "",log="y",
	ylab = "Number of resistance occurences",col=c("black","grey"),
	ylim=c(5,500),las=3,font=3,yaxt="n")
axis(2)
mtext("Country", side=1, line=6)
legend("topleft",legend=c("Before 2022","2022-2031"),fill=c("black","grey"))


barplot(prop.country, xlab = "",
	ylab = "Proportion of tested occurences predicted to evolve resistance",
	las=3,font=3,yaxt="n")
axis(2)
mtext("Country", side=1, line=6)

dev.off()


########
###
###Future resistance in species
###
########


future.spec<-table(future.resist$Species)
past.spec<-table(ML.data$Species[
	which(ML.data$Resistance=="A")])
potential.spec<-table(test$Species)


future.spec[names(past.spec)[which(names(past.spec)%in%
	names(future.spec)==F)]]<-0
future.spec<-future.spec[names(past.spec)]


potential.spec[names(past.spec)[which(names(past.spec)%in%
	names(potential.spec)==F)]]<-0
potential.spec<-potential.spec[names(past.spec)]


sort(future.spec/potential.spec)
sort(past.spec)
write.csv(cbind(past.spec,future.spec,potential.spec,future.spec/potential.spec),
	file="species_test.csv")


combined<-rbind(past.spec,future.spec)
colnames(combined)<-names(past.spec)
rownames(combined)<-c("Past","Predicted")

prop<-as.vector(future.spec/potential.spec)
names(prop)<-names(future.spec)


specs.to.plot<-c("Lolium rigidum","Lolium perenne ssp. multiflorum",
	"Conyza canadensis","Eleusine indica","Lactuca serriola",
	"Alopecurus myosuroides","Amaranthus palmeri","Amaranthus retroflexus",
	"Avena fatua")

specs.to.plot<-c("Lolium rigidum","Lolium perenne ssp. multiflorum",
	"Kochia scoparia","Amaranthus palmeri","Avena fatua")

specs.to.plot<-c("Lolium rigidum","Lolium perenne ssp. multiflorum",
	"Conyza canadensis","Eleusine indica",
	"Alopecurus myosuroides","Amaranthus palmeri","Amaranthus retroflexus",
	"Avena fatua")

tiff(filename = "Figure S5_analysis3.tif",width = 20, height = 20, 
	units = "cm",res=300, compression = "lzw")
par(mfrow=c(1,2))
par(mar=c(15,5,1,1))
barplot(combined[,specs.to.plot], xlab = "",log="y",
	ylab = "Number of resistance occurences",col=c("black","grey"),
	ylim=c(5,7500),las=3,font=3,yaxt="n")
axis(2)
mtext("Species", side=1, line=12)
legend("topleft",legend=c("Before 2021","2021-2031"),fill=c("black","grey"))


barplot(prop[specs.to.plot], xlab = "",
	ylab = "Proportion of tested occurences predicted to evolve resistance",
	las=3,font=3,yaxt="n")
axis(2)
mtext("Species", side=1, line=12)


dev.off()


#####
##
## Plot maps of resistance change in species
##
#####


CABI.weed<-read.csv(file.choose(),header=F)


plot.spec<-"Lactuca serriola"


spec.countries<-CABI.weed[which(CABI.weed[,2]==plot.spec),1]
unique(test.weed$Country[which(test.weed$Species==plot.spec)])%in%spec.countries

past.spec.resist<-array(dim=length(spec.countries),data=0)
names(past.spec.resist)<-spec.countries

past.spec.resist[names(table(ML.data$Country[
	which(ML.data$Resistance=="A" & ML.data$Species==plot.spec)]))]<-
	table(ML.data$Country[
	which(ML.data$Resistance=="A" & ML.data$Species==plot.spec)])


future.spec.resist<-array(dim=length(spec.countries),data=0)
names(future.spec.resist)<-spec.countries

future.spec.resist[names(table(future.resist$Country[
	which(future.resist$Species==plot.spec)]))]<-
	table(future.resist$Country[which(future.resist$Species==plot.spec)])


potential.spec.resist<-array(dim=length(spec.countries),data=0)
names(potential.spec.resist)<-spec.countries

potential.spec.resist[names(table(test$Country[
	which(test$Species==plot.spec)]))]<-
	table(test$Country[which(test$Species==plot.spec)])


future.spec.percent<-future.spec.resist/potential.spec.resist
future.spec.percent[is.na(future.spec.percent)]<-0
future.spec.percent[which(future.spec.percent==Inf)]<-future.spec.resist[which(future.spec.percent==Inf)]

future.spec.resist


names(future.spec.percent)[
	names(future.spec.percent)%in%countryExData$Country==F]


names(future.spec.resist)[
	names(future.spec.resist)%in%countryExData$Country==F]


names(past.spec.resist)[
	names(past.spec.resist)%in%countryExData$Country==F]


countryExData$Country[15]<-"Bosnia and Herzegovina"
countryExData$Country[21]<-"Central African Republic"
countryExData$Country[26]<-"C�te D'ivoire"
countryExData$Country[28]<-"Democratic Republic of Congo"
countryExData$Country[34]<-"Czech Republic"
countryExData$Country[38]<-"Dominican Republic"
countryExData$Country[145]<-"Vietnam"
countryExData$Country[134]<-"Trinidad and Tobago"


past.spec.resist<-past.spec.resist[names(past.spec.resist)%in%countryExData$Country]
future.spec.percent<-future.spec.percent[names(future.spec.percent)%in%countryExData$Country]
future.spec.resist<-future.spec.resist[names(future.spec.resist)%in%countryExData$Country]


country.codes<-countryExData[which(countryExData$Country%in%names(past.spec.resist)),
	c("Country","ISO3V10")]

uncertain<-array(dim=length(spec.countries),data=0)
names(uncertain)<-spec.countries
uncertain[which(spec.countries%in%ML.data$Country==F)]<-1


plot.past.spec.country<-data.frame(as.array(past.spec.resist[country.codes[,1]]+1),
	country.codes,uncertain[country.codes[,1]])
names(plot.past.spec.country)<-c("Freq","Name","Code","Data Available")


past.spec.map <- joinCountryData2Map( plot.past.spec.country
	,joinCode = "ISO3"
	,nameJoinColumn = "Code")


plot.futurecount.spec.country<-data.frame(as.array(future.spec.resist[country.codes[,1]]+1),
	country.codes,uncertain[country.codes[,1]])
names(plot.futurecount.spec.country)<-c("Freq","Name","Code","Data Available")


futurecount.spec.map <- joinCountryData2Map( plot.futurecount.spec.country
	,joinCode = "ISO3"
	,nameJoinColumn = "Code")


plot.future.spec.country<-data.frame(as.array(future.spec.percent[country.codes[,1]]),
	country.codes,uncertain[country.codes[,1]])
names(plot.future.spec.country)<-c("Freq","Name","Code","Data Available")


future.spec.map <- joinCountryData2Map( plot.future.spec.country
	,joinCode = "ISO3"
	,nameJoinColumn = "Code")


tiff(filename = "Lactuca.tif",width = 20, height = 10, 
	units = "cm",res=300, compression = "lzw")

par(mar=c(0,1,0,0.5))
par(mfrow=c(1,3))
mapCountryData(past.spec.map,nameColumnToPlot="Freq",
	catMethod="logFixedWidth",colourPalette=c("blue","grey","red"),
	borderCol="black",numCats = 10,nameColumnToHatch="Data Available",
	 mapTitle="")


mapCountryData(futurecount.spec.map,nameColumnToPlot="Freq",
	catMethod="logFixedWidth",colourPalette=c("blue","grey","red"),
	borderCol="black",numCats = 10,nameColumnToHatch="Data Available",
	 mapTitle="")


mapCountryData(future.spec.map,nameColumnToPlot="Freq",
	catMethod="fixedWidth",colourPalette=c("blue","grey","red"),
	borderCol="black",numCats = 10,nameColumnToHatch="Data Available",
	 mapTitle="")

dev.off()


#####
##
## resistance change in situation
##
#####

future.sit<-table(future.resist$Situations)
past.sit<-table(ML.data$Situations[which(ML.data$Resistance=="A")])
potential.sit<-table(test$Situations)

future.sit[names(past.sit)[which(names(past.sit)%in%
	names(future.sit)==F)]]<-0
future.sit<-future.sit[names(past.sit)]

potential.sit[names(past.sit)[which(names(past.sit)%in%
	names(potential.sit)==F)]]<-0
potential.sit<-potential.sit[names(past.sit)]


sort(future.sit/potential.sit)
sort(past.sit)


write.csv(cbind(past.sit,future.sit,potential.sit,future.sit/potential.sit),
	file="sit_test.csv")


combined<-rbind(past.sit,future.sit)
colnames(combined)<-names(future.sit)
rownames(combined)<-c("Past","Predicted")

prop<-as.vector(future.sit/potential.sit)
names(prop)<-names(future.sit)


sits.to.plot<-c("Wheat","Soybean","Corn (maize)","Rice","Roadsides",
	"Spring Barley","Grapes")

sits.to.plot<-c("Cereals","Beans","Corn (maize)","Roadsides",
	"Fruit")

sits.to.plot<-c("Cereals","Beans","Corn (maize)", "Rice","Roadsides",
	"Fruit")

tiff(filename = "Figure S6_analysis3.tif",width = 20, height = 20, 
	units = "cm",res=300, compression = "lzw")
par(mfrow=c(1,2))
par(mar=c(8,5,1,1))
barplot(combined[,sits.to.plot], xlab = "",log="y",
	ylab = "Number of resistance occurences",col=c("black","grey"),
	,las=3,font=3,yaxt="n")
axis(2)
mtext("Situation", side=1, line=7)
legend("topright",legend=c("Before 2021","2021-2031"),fill=c("black","grey"))


barplot(prop[sits.to.plot], xlab = "",
	ylab = "Proportion of tested occurences predicted to evolve resistance",
	las=3,font=3,yaxt="n")
axis(2)
mtext("Situation", side=1, line=7)


dev.off()


#####
##
## Plot maps of resistance change in situations
##
#####


trade.crop<-read.csv(file.choose(),header=T)


plot.sit<-"Grapes"


sit.countries<-strsplit(trade.crop[which(trade.crop[,1]==plot.sit),2],split=", ")[[1]]
sit.countries<-c(sit.countries,
	unique(test.weed$Country[which(test.weed$Situations==plot.sit)])[
	which(unique(test.weed$Country[which(test.weed$Situations==plot.sit)])%in%sit.countries==F)])


for(i in 1:nrow(crop.categories))
{
	if(crop.categories[i,2]==plot.sit)
	{
		
		sit.countries<-c(sit.countries,
			strsplit(trade.crop[which(trade.crop[,1]==crop.categories[i,1]),2],split=", ")[[1]])

		sit.countries<-c(sit.countries,
			unique(test.weed$Country[which(test.weed$Situations==crop.categories[i,1])])[
			which(unique(test.weed$Country[which(test.weed$Situations==crop.categories[i,1])])%in%sit.countries==F)])

		plot.sit<-c(plot.sit,crop.categories[i,1])
	}
}


past.sit.resist<-array(dim=length(sit.countries),data=0)
names(past.sit.resist)<-sit.countries

past.sit.resist[names(table(ML.data$Country[
	which(ML.data$Resistance=="A" & ML.data$Situations%in%plot.sit)]))]<-
	table(ML.data$Country[
	which(ML.data$Resistance=="A" & ML.data$Situations%in%plot.sit)])


future.sit.resist<-array(dim=length(sit.countries),data=0)
names(future.sit.resist)<-sit.countries

future.sit.resist[names(table(future.resist$Country[
	which(future.resist$Situations%in%plot.sit)]))]<-
	table(future.resist$Country[which(future.resist$Situations%in%plot.sit)])


potential.sit.resist<-array(dim=length(sit.countries),data=0)
names(potential.sit.resist)<-sit.countries

potential.sit.resist[names(table(test$Country[
	which(test$Situations%in%plot.sit)]))]<-
	table(test$Country[which(test$Situations%in%plot.sit)])


future.sit.percent<-future.sit.resist/potential.sit.resist
future.sit.percent[is.na(future.sit.percent)]<-0
future.sit.percent[which(future.sit.percent==Inf)]<-future.sit.resist[which(future.sit.percent==Inf)]


names(future.sit.resist)[
	names(future.sit.resist)%in%countryExData$Country==F]


names(future.sit.percent)[
	names(future.sit.percent)%in%countryExData$Country==F]


names(past.sit.resist)[
	names(past.sit.resist)%in%countryExData$Country==F]

countryExData$Country[15]<-"Bosnia and Herzegovina"
countryExData$Country[21]<-"Central African Republic"
countryExData$Country[26]<-"C�te D'ivoire"
countryExData$Country[28]<-"Democratic Republic of Congo"
countryExData$Country[34]<-"Czech Republic"
countryExData$Country[38]<-"Dominican Republic"
countryExData$Country[145]<-"Vietnam"
countryExData$Country[134]<-"Trinidad and Tobago"


past.sit.resist<-past.sit.resist[names(past.sit.resist)%in%countryExData$Country]
future.sit.resist<-future.sit.resist[names(future.sit.resist)%in%countryExData$Country]
future.sit.percent<-future.sit.percent[names(future.sit.percent)%in%countryExData$Country]


country.codes<-countryExData[which(countryExData$Country%in%names(past.sit.resist)),
	c("Country","ISO3V10")]

uncertain<-array(dim=length(sit.countries),data=0)
names(uncertain)<-sit.countries
uncertain[which(sit.countries%in%ML.data$Country==F)]<-1


plot.past.sit.country<-data.frame(as.array(past.sit.resist[country.codes[,1]]+1),
	country.codes,uncertain[country.codes[,1]])
names(plot.past.sit.country)<-c("Freq","Name","Code","Data Available")


past.sit.map <- joinCountryData2Map( plot.past.sit.country
	,joinCode = "ISO3"
	,nameJoinColumn = "Code")


plot.futurecount.sit.country<-data.frame(as.array(future.sit.resist[country.codes[,1]]+1),
	country.codes,uncertain[country.codes[,1]])
names(plot.futurecount.sit.country)<-c("Freq","Name","Code","Data Available")


futurecount.sit.map <- joinCountryData2Map( plot.futurecount.sit.country
	,joinCode = "ISO3"
	,nameJoinColumn = "Code")


plot.future.sit.country<-data.frame(as.array(future.sit.percent[country.codes[,1]]),
	country.codes,uncertain[country.codes[,1]])
names(plot.future.sit.country)<-c("Freq","Name","Code","Data Available")


future.sit.map <- joinCountryData2Map( plot.future.sit.country
	,joinCode = "ISO3"
	,nameJoinColumn = "Code")


tiff(filename = "spring barley.tif",width = 20, height = 10, 
	units = "cm",res=300, compression = "lzw")


par(mar=c(0,1,0,0.5),mfrow=c(1,3))
mapCountryData(past.sit.map,nameColumnToPlot="Freq",
	catMethod="logFixedWidth",colourPalette=c("blue","grey","red"),
	borderCol="black",numCats = 10,nameColumnToHatch="Data Available",
	 mapTitle="")
mapCountryData(futurecount.sit.map,nameColumnToPlot="Freq",
	catMethod="logFixedWidth",colourPalette=c("blue","grey","red"),
	borderCol="black",numCats = 10,nameColumnToHatch="Data Available",
	 mapTitle="")


mapCountryData(future.sit.map,nameColumnToPlot="Freq",
	catMethod="fixedWidth",colourPalette=c("blue","grey","red"),
	borderCol="black",numCats = 10,nameColumnToHatch="Data Available",
	 mapTitle="")


dev.off()


#####
##
## resistance change in Actives
##
#####

future.act<-table(future.resist$Actives)
past.act<-table(ML.data$Actives[which(ML.data$Resistance=="A")])
potential.act<-table(test$Actives)

future.act[names(past.act)[which(names(past.act)%in%
	names(future.act)==F)]]<-0
future.act<-future.act[names(past.act)]


potential.act[names(past.act)[which(names(past.act)%in%
	names(potential.act)==F)]]<-0
potential.act<-potential.act[names(past.act)]


sort(future.act/potential.act)
sort(past.act)


write.csv(cbind(past.act,future.act,potential.act,future.act/potential.act),
	file="act_test.csv")


#####
##
## resistance change by MOA
##
#####
MOAs<-read.csv(file.choose(),header=F,row.name=1)

future.MOA<-cbind(future.act,MOAs[names(future.act),])
future.MOA<-aggregate(future.MOA[,1], by=list(future.MOA[,2]), FUN=sum)

past.MOA<-cbind(past.act,MOAs[names(past.act),])
past.MOA<-aggregate(past.MOA[,1], by=list(past.MOA[,2]), FUN=sum)

potential.MOA<-cbind(potential.act,MOAs[names(potential.act),])
potential.MOA<-aggregate(potential.MOA[,1], by=list(potential.MOA[,2]), FUN=sum)


percent.inc.MOA<-future.MOA[,2]/potential.MOA[,2]
names(percent.inc.MOA)<-future.MOA[,1]
sort(percent.inc.MOA)

barplot(percent.inc.MOA)


combined<-rbind(past.MOA[,2],future.MOA[,2])
colnames(combined)<-future.MOA[,1]
rownames(combined)<-c("Past","Predicted")


tiff(filename = "Figure S7analysis3.tif",width = 20, height = 25, 
	units = "cm",res=300, compression = "lzw")

par(mfrow=c(2,1))

barplot(combined, xlab = "MOA", ylab = "Number of resistance occurences",
	,col=c("black","grey"),log="y")
legend("topright",legend=c("Before 2021","2021-2031"),fill=c("black","grey"))
barplot(percent.inc.MOA,ylab="Proportion of tested occurences predicted to evolve resistance",
	xlab="MOA")

dev.off()
```
