## Supplementary Methods for "Predicting global herbicide resistance hotspots using a 30-year-old database and machine-learning techniques": Supplementary Data 30.R

```
resist.data<-read.csv(file.choose())#####Read in supplementary data 1 (the original database downloaded)


countries<-unique(resist.data$Country)######unique countries

country.res<-vector(length=length(countries))######vector to store number of resistance cases in each country
country.u<-vector(length=length(countries))#######vector to store good's u values


for(i in 1:length(countries))#####each country 1 by 1
{
	country.occ<-resist.data[resist.data$Country==countries[i],]#######cases from that country
	country.obs<-nrow(country.occ)#####number of cases observed
	country.sing<-length(which(table(country.occ$Species)==1))#####number of singletons

	country.res[i]<-country.obs#####store nummber of cases
	country.u[i]<- 1-(country.sing/country.obs)####store good's u
}

names(country.u)<-countries
names(country.res)<-countries


cor.test(country.u,log10(country.res),method="pearson")#####correlation test


#######number of researchers

data<-read.csv(file.choose(),row.name=1)####Read in supplementary data 11 (count of researchers derived from the database)


cor.test(log10(data[intersect(which(data[,1]>0),####correlartuon test (with 0 values removed to allow log transformation)
	which(data[,2]>0)),1]),log10(data[intersect(
	which(data[,1]>0),which(data[,2]>0)),2]),method="pearson")
```
